# A BTB-Domain Transcription factor Recruits Chromatin Remodelers and a Histone Chaperone during the exit from Pluripotency

**DOI:** 10.1101/2020.12.03.409912

**Authors:** Daniel Olivieri, Sujani Paramanathan, Anaïs F. Bardet, Daniel Hess, Sébastien A. Smallwood, Ulrich Elling, Joerg Betschinger

## Abstract

Transcription factors (TFs) harboring a btb (Broad-Complex, Tramtrack and Bric a brac) domain play important roles in development and disease. They are thought to recruit transcriptional modulators to DNA through their btb domain. However, a systematic molecular understanding of this TF family is lacking. Here, we identify the zinc finger btb-TF *Zbtb2* in a genetic screen for regulators of exit from pluripotency and dissect its mechanistic mode of action. We show that ZBTB2 binds the chromatin remodeler Ep400 to mediate downstream transcription. Independently, the btb domain directly interacts with the chromatin remodeller NuRD and the histone chaperone HiRA via the GATAD2A/B and UBN2 subunits, respectively. NuRD recruitment is a common feature of btb-TFs and we propose by phylogenetic analysis that this is an evolutionary ancient property. Binding to UBN2, in contrast, is specific to ZBTB2 and requires a C-terminal extension of the btb domain. This study therefore identifies a btb-domain TF that recruits chromatin modifiers and a histone chaperone during a developmental cell state transition, and defines unique and shared molecular functions of the btb-domain TF family.

## INTRODUCTION

Transcription factors (TFs) are key determinants of gene expression and, therefore, play a major role in development and disease (Lambert et al., 2018). TFs interpret the regulatory code of the genome by binding to DNA and regulating transcription (Lambert et al., 2018). While DNA-binding is well characterized (Weirauch et al., 2014), the mechanisms by which TFs modulate transcription are not completely understood. Many TFs present a modular protein architecture containing DNA-binding domains and domains that interact with transcriptional activators or repressors (Lambert et al., 2018). Zinc-finger domains (Znfs) are the most common family of DNA-binding domains and are often found in combination with Krueppel associated box (KRAB) or Broad-Complex, Tramtrack and Bric a brac (btb) domains (Collins et al., 2001). KRAB domains recruit KAP1 (Helleboid et al., 2019) and therefore mediate transcriptional repression. In contrast, there is no comprehensive understanding of the transcriptional role of btb domains.

There are three groups of TFs containing btb domains (btb-TFs): the *Zbtb*, the *Bach*, and the *Nacc* families, which are defined by their DNA binding domains: Znf, bZIP, and BEN, respectively (Stogios et al., 2005). In human and mouse, the *Bach* and the *Nacc* families contain only 2 members each, and the *Zbtb* family comprises 49 members. Several of them are critical regulators of fate allocation and differentiation across many organs and systems (Chevrier and Corcoran, 2014). A striking example is hematopoiesis, in which btb-TFs direct the differentiation of several lineages (Maeda, 2016). The btb domains are invariably found at the N-terminus of btb-TFs and the DNA-binding domains at the C-terminus, separated by a long non-conserved linker region (Maeda, 2016). Although btb domains function in homo- and heterodimerization and protein-protein interactions, the btb domains found in TFs and CUL3 ubiquitin ligases constitute separate families (Stogios et al., 2005). Mechanistic studies of Zbtb factors have focused on the proto-oncogenes *Bcl6* (*Zbtb27*) and *Lrf* (*Zbtb7a*) and on the tumor suppressor *Plzf* (*Zbtb16*), and found that they act as transcriptional repressors via the recruitment of complexes such as NCOR/SMRT, BCOR, SIN3A/B, and NuRD (Maeda, 2016).

Mouse embryonic stem cells (mESCs) are a developmentally relevant cell type that captures the pluripotent state of the pre-implantation mouse epiblast, and that recapitulates developmental progression upon release into differentiation *in vitro* (Martello and Smith, 2014). Furthermore, a large number of genomic datasets have been derived from mESCs, making them an ideal model for mechanistic studies of transcription. mESCs are maintained by provision of the cytokine leukemia inhibitory factor (LIF) in a fetal calf serum containing medium (Serum-LIF) (Smith et al., 1988) or of inhibitors (i) of glycogen synthetase kinase 3 (GSK3) and mitogen-activated protein kinase kinase (MEK) in a chemically defined medium (N2B27) (Ying et al., 2008). LIF, GSK3(i) and MEK(i) stabilize a pluripotency TF network centered on OCT4, SOX2 and NANOG (Martello and Smith, 2014). mESCs grown in the presence of the two inhibitors (2i) are called naïve and display higher and more homogeneous expression levels of pluripotency TFs than Serum-LIF grown cells (Silva and Smith, 2008). Naïve and Serum-LIF cell states are interconvertible (Galonska et al., 2015), while naïve cells can efficiently differentiate into post-implantation epiblast-like cells (EpiLCs) (Hayashi et al., 2011). In both naïve and Serum-LIF conditions, mESCs transition through a 2-cell-embryo (2C)-like state that is thought to reflect properties of cleavage-stage embryos, and that is characterized by expanded developmental potency (Macfarlan et al., 2012), increased histone mobility (Bošković et al., 2014) and a specific gene expression signature which includes the upregulation of endogenous retroviruses (Macfarlan et al., 2011). While the TF *Dux* directly binds and activates the promoters of 2C-like genes (Hendrickson et al., 2017), it is less clear how other chromatin regulators, such as PRC1.6, Ep400 (Rodriguez-Terrones et al., 2018), or CAF-1 (Ishiuchi et al., 2015) mechanistically regulate 2C-like genes.

The Nucleosome Remodeling and Deacetylase (NuRD) complex is composed of HDAC1/2, CHD4, GATAD2A/B, RBBP4/7, MTA1/2/3, and MBD2/3 (Xue et al., 1998). MBD2 and MBD3 are mutually exclusive subunits and define the two functionally distinct MBD2-NuRD and MBD3-NuRD complexes (Guezennec et al., 2006). *Mbd3* has been subject to several genetic studies in mESCs, showing that NuRD drives mESC differentiation (Kaji et al., 2007, 2006; Reynolds et al., 2012) and fine-tunes gene expression by modulating chromatin accessibility (Bornelöv et al., 2018).

The Histone Regulator A (HiRA) complex acts as the H3.3 histone chaperone at euchromatic loci (Goldberg et al., 2010). HiRA has been proposed to be recruited by naked DNA and to have a nucleosome-gap filling function (Ray-Gallet et al., 2011). It is composed of the subunits HIRA, CABIN1, and UBN1/2 (Tagami et al., 2004). It has been suggested that UBN1 and UBN2 are part of two independent, but functionally indistinguishable, complexes, UBN1-HiRA and UBN2-HiRA (Xiong et al., 2018). Euchromatic H3.3 is found in H2A.Z containing nucleosomes (Jin and Felsenfeld, 2007) that are incorporated into chromatin by the chromatin remodeling complex Ep400 (Pradhan et al., 2016), yet if and how Ep400 interacts with H3.3 chaperones is unclear. HiRA is required for exit from the naïve pluripotency during differentiation (Leeb et al., 2014), while the Ep400 complex is essential for the maintenance of mESCs (Fazzio et al., 2008).

Here, we identify the btb-TF *Zbtb2* in a genetic screen for regulators of exit from mESC pluripotency, and report a detailed mechanistic analysis of its function, showing that ZBTB2 recruits ZNF639, MBD3-NuRD, UBN2-HiRA, and the Ep400 complex. Transcriptome analysis reveals that ZBTB*2* interactors form two functionally distinct modules, one encompassing ZNF639, NuRD and HiRA, and the other corresponding to the Ep400 complex. We show that NuRD and HiRA associate with ZBTB2 via the subunits GATAD2A/B and UBN2, respectively. We systematically test these interactions across all btb-TFs in yeast-2-hybrid (Y2H) screens. We find that ZBTB2 harbors an extension of the btb domain that mediates a unique interaction with UBN2. GATAD2A/B interaction is instead a common feature of btb-TFs and shared across several btb-TF phylogenetic branches, making NuRD recruitment a candidate ancestral feature of TF-associated btb domains. Our study therefore reveals unique and shared molecular functions of the btb-domain TF family.

## RESULTS

### A sensitized genetic screen identifies Zbtb2 as regulator of the exit from pluripotency

We performed a sensitized genetic screen for maintenance of pluripotency (Ying et al., 2003) in the presence of the GSK3 inhibitor CHIR99021 (CHIR) (Sato et al., 2004), which is unable to block mESC differentiation in the absence of LIF or Mek(i) (Wray et al., 2010). As the role of CHIR in mESCs maintenance is well characterized (Martello et al., 2012; Wray et al., 2011), this medium formulation should increase sensitivity for other, less understood, pathways. We mutagenized haploid mESCs (Elling et al., 2011) harboring an *Oct4>GFP-puro* reporter with retroviruses carrying a splicing acceptor site for insertional mutagenesis and Oct4 enhancer elements for overexpression (Schnütgen et al., 2008), leading to both loss and gain of function alleles. We cultured the mutagenized library in N2B27 +CHIR +puromycin to select for undifferentiated cells and harvested the cells after 16 to 23 days. We mapped the insertions by high-throughput inverse PCR and determined insertion enrichment compared to the starting library for every gene (Fig. 1A, Table S1). Confirming the specificity of our setup, *Fgfr1* and *Lif* were among the highest scoring hits. FGFR1 is the main FGF receptor in mESCs and acts upstream of MEK activation (Molotkov et al., 2017), and chemical inhibition of FGFR signaling has been shown to substitute for MEK(i) in mESC maintenance (Ying et al., 2008). *Lif*, evidently a gain of function hit, is also able to sustain pluripotency in combination with CHIR (Sato et al., 2004). *Oct4* is a technical false positive hit, as it can drive the expression of the *Oct4>GFP-puro* reporter irrespectively of the cell state. Confidence in our analysis was also bolstered by the identification of *Esrrb* and *Tfcp2l1* amongst the insertion-depleted genes, as these are transcriptional mediators of CHIR activity (Martello et al., 2013, 2012; Qiu et al., 2015). Some of the highest scoring screen hits, such as *Cbx1* (Mattout et al., 2015), *Eed* (Leeb et al., 2010; Tee et al., 2014), *Trp53* (Lin et al., 2005) and *Upf2* (Li et al., 2015), have previously been implicated in the exit from pluripotency. We therefore decided to validate the hits *Zbtb2, Zfp42, Nexmif*, and *Nmt1*.

**Figure 1:**
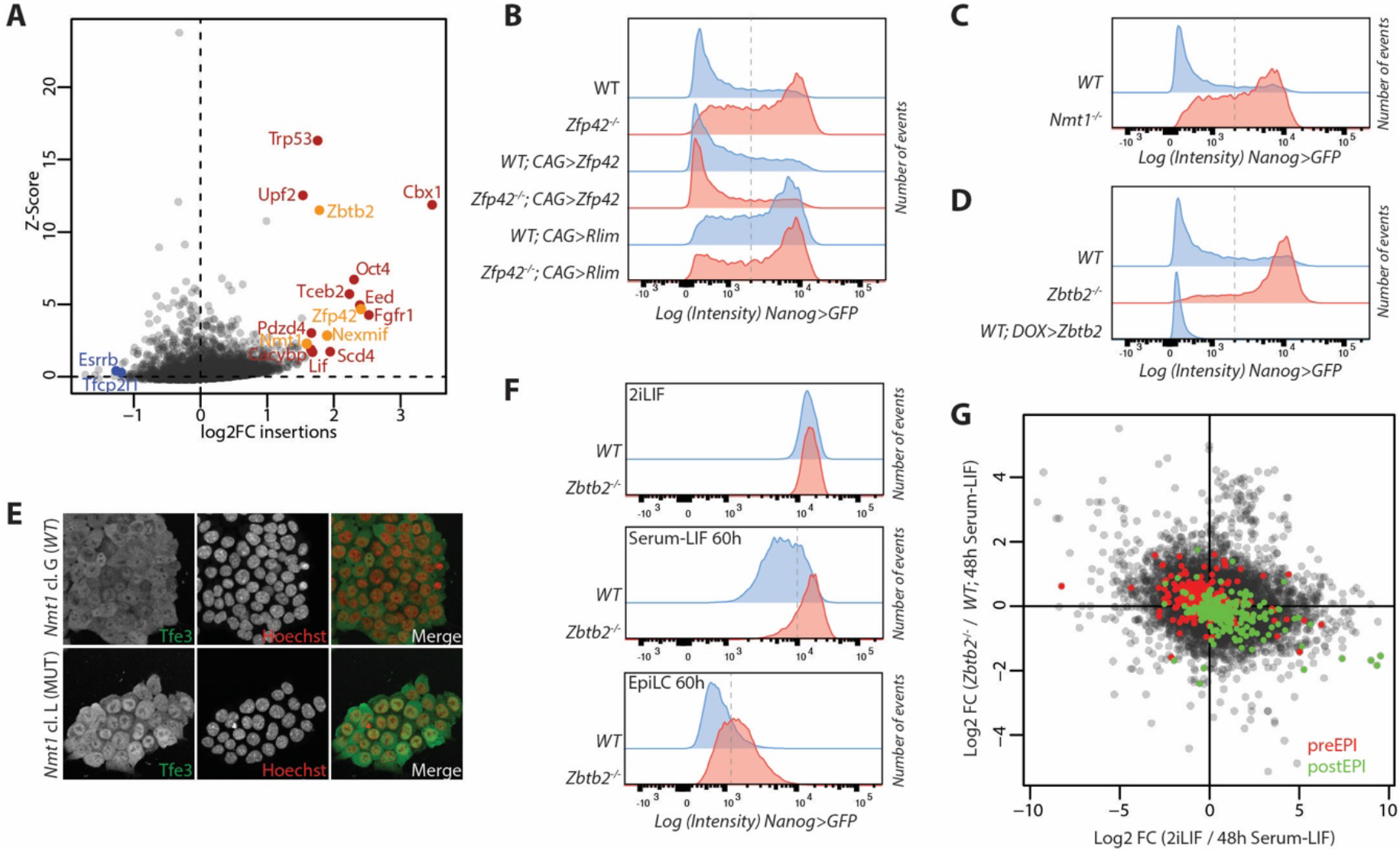
Screen results and validation. **(A**) Z-scores and enrichment fold-changes of retroviral insertions within the gene bodies of indicated genes **(B-D**,**F)** *Nanog>GFP* intensities after 3 days in N2B27 +CHIR +bFGF (**B-D**) and as indicated (**F**) of indicated genotypes and treatments. Dashed lines indicate the thresholds for quantifications presented in Fig. S1D,F. **(E)** Tfe3 immunofluorescence in *Nmt1*^-/-^ and WT mESCs in 2iLIF. Nuclei were counterstained with Hoechst. **(G)** Scatterplot of log2 fold-changes (FC) in gene expression of indicated contrasts. Green labels post-implantation and red pre-implantation epiblast specific genes (Boroviak et al., 2015).

The *Nexmif* gene lies upstream of the *Rlim* transcription start site and RLIM is a known E3 ubiquitin ligase for ZFP42 (Gontan et al., 2012), so we reasoned that the *Nexmif* insertion enrichment would lead to *Rlim* overexpression. We therefore generated CRISPR knock-out clones (Table S2) for *Zfp42* (Fig. S1A), *Nmt1* (Fig. S1B) and *Zbtb2* (Fig. S1C), and *Rlim* overexpressing cells using naïve TNG-A mESCs, a conventional diploid cell line harboring a *Nanog>GFP* reporter (Chambers et al., 2007). Upon exposure to N2B27 +CHIR, concomitant with the addition of recombinant basic FGF (bFGF) to increase the stringency of the assay, all three mutants and the *Rlim* overexpressing cells showed delayed down-regulation of the *Nanog* reporter when compared to wildtype (*WT*) cells (Fig. 1B, C, D, and Fig. S1D), indicating that they are *bona fide* loss of function screen hits. Overexpression of *Rlim* in the *Zfp42* mutant did not modify the phenotype of the *Zfp42* single mutant (Fig. 1B and Fig. S1D), supporting the epistatic relationship between *Rlim* and *Zfp42*. While *Zfp42* overexpression did not affect differentiation (Fig. 1B and Fig. S1D), strong *Zbtb2* constitutive overexpression using a CAG promoter caused cell death (not shown). We therefore tested the effect of moderate *Zbtb2* overexpression, achieved by a doxycycline (DOX) inducible promoter, and observed accelerated differentiation upon *Zbtb2* induction (Fig. 1D and Fig. S1D).

*Nmt1* encodes an N-myristoyltransferase (Yang et al., 2005) and myristoylation is required for the function of FRS2, an essential component for FGFR1 signaling (Kouhara et al., 1997). We therefore speculated that loss of *Nmt1* would inhibit mESC differentiation by dampening mitogen activated protein kinase (MAPK) signaling. However, ERK phosphorylation upon exposure to bFGF was unperturbed in *Nmt1*^-/-^ cells (Fig. S1E). We therefore turned our attention to another myristoylated protein, LAMTOR1 (Thinon et al., 2014), which is required for TFE3 nuclear exclusion (Villegas et al., 2019) and, in turn, for the exit from pluripotency (Betschinger et al., 2013). Immunofluorescence staining revealed abnormal nuclear localization of TFE3 in *Nmt1*^-/-^ mESCs (Fig. 1E), suggesting that ectopically active TFE3 mediates the differentiation delay in the absence of *Nmt1*.

Although *Zbtb2* had already been suggested to play a role in the differentiation of Serum-LIF mESCs (Karemaker and Vermeulen, 2018), we decided to focus our efforts on this factor in the hope of gaining insights that are broadly applicable to btb-TFs. We first sought to better characterize *Zbtb2*’s role in the exit from the naïve state and determine when the earliest developmental defect would arise. We first tested differentiation of 2iLIF cells using the EpiLC differentiation protocol, which faithfully mimics the pre- to post-implantation epiblast transition (Hayashi et al., 2011), and observed a delay in *Nanog* reporter downregulation in *Zbtb2*^-/-^ cells although *Nanog* levels were unchanged in steady-state 2iLIF cultures (Fig. 1F and Fig.S1F). We then turned to differentiation in Serum-LIF, which establishes a developmentally advanced pluripotent state (Kalkan and Smith, 2014). Even in this assay, *Nanog>GFP* downregulation was delayed in *Zbtb2*^-/-^ cells (Fig. 1F and Fig.S1F). To determine transcriptome-wide changes we performed RNA sequencing (RNAseq) of *WT* and *Zbtb2*^-/-^ cells in 2iLIF and during differentiation. We found that gene expression changes that accompany the 2iLIF to Serum-LIF transition in *WT* cells were dampened in *Zbtb2*^-/-^cells exposed to Serum-LIF for 48 hours (h) (Pearson correlation coefficient R = −0.19, Fig. 1G, Table S3). When we specifically focused on genes regulated during embryonic development (Boroviak et al., 2015), we found that *Zbtb2*^-/-^ cells in Serum-LIF, when compared to *WT* controls, were impaired in upregulating genes that are expressed in the post-implantation epiblast and in downregulating genes that are predominantly transcribed in the pre-implantation epiblast (Fig. 1G). This coherent deregulation of developmental genes was specific to the Serum-LIF transition and undetectable in steady-state 2iLIF cells (Fig. S1G). In summary, loss of *Zbtb2* delays and *Zbtb2* overexpression increases mESC differentiation, demonstrating an instructive role of ZBTB2 in cell state transitions.

### An extended btb domain binds UBN2 and GATAD2B; NuRD interaction is stabilized by ZNF639

To understand the mechanisms by which *Zbtb2* exerts its function, we performed affinity purification–mass spectrometry (AP-MS) of ZBTB2 in mESCs. In the absence of antibodies detecting the endogenous protein we generated mESCs expressing an avidin (AVI)-tagged *Zbtb2* transgene which, similar to untagged *Zbtb2*, induces differentiation when overexpressed (Fig. S2A), therefore confirming functionality. Using streptavidin pull-downs, we identified ZBTB25 and ZNF639, previously reported to interact with ZBTB2 (Karemaker and Vermeulen, 2018), and all subunits of the NuRD and of the HiRA complexes as specific ZBTB2 interactors (Fig. 2A). We did not detect MBD2 or UBN1 peptides, demonstrating co-purification of MBD3-NuRD and UBN2-HiRA complexes, specifically. To better understand how ZBTB2 recruits such a complex interactome, we performed AP-MS of ZBTB2-AVI alleles harboring mutations in the btb and Znf domains (Table S4). All experiments were carried out in a *Zbtb2*^-/-^*Zbtb25*^-/-^ background (Fig. S2B) to prevent indirect interactions due to bait dimerization with endogenous ZBTB2 or ZBTB25, and in the presence of Benzonase nuclease to avoid DNA- and RNA-bridged interactions. Mutation of the btb domain caused loss of the interaction with UBN2-HiRA and of Znf1 abolished the interaction with ZNF639 and NuRD, while mutation of the other Znfs did not significantly alter the interactome (Fig. 2B, Table S5). However, loss of interaction upon domain mutation does not imply direct physical binding, as it could be due to an indirect functional dependency or bridging factor.

**Figure 2:**
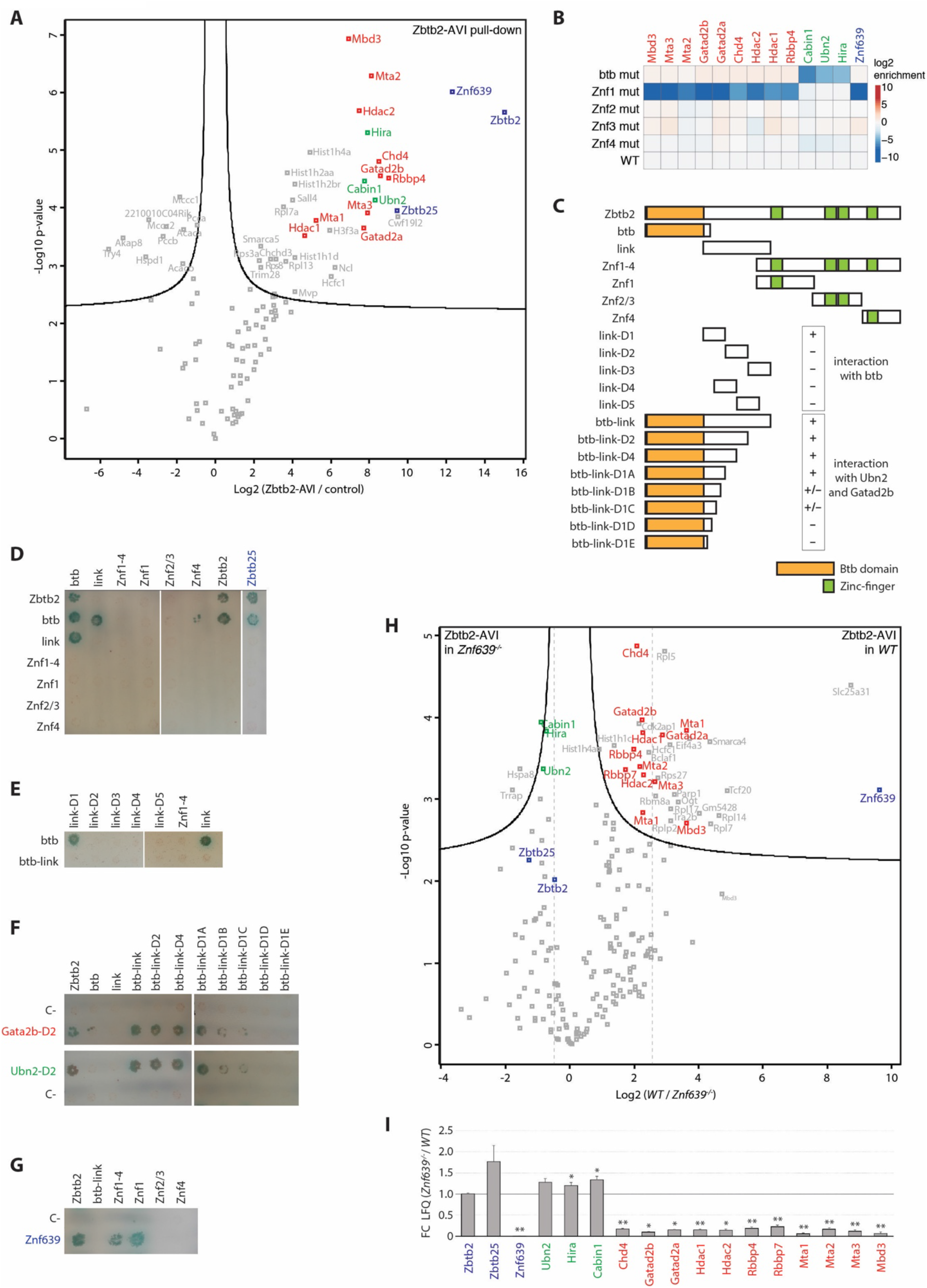
An extended btb domain binds UBN2 and GATAD2B; NuRD interaction is stabilized by ZNF639. **(A)** Volcano plot of protein enrichments in AP-MS of ZBTB2-AVI compared to control BirA-expressing cells in 2iLIF. ZBTB2 and partner TFs are indicated in blue, NuRD subunits in red, and HiRA subunits in green. **(B)** Bait-normalized log2 fold enrichment of ZNF639, HiRA subunits and NuRD subunits in AP-MS of ZBTB2-AVI alleles with indicated mutation compared to wildtype ZBTB2-AVI in *Zbtb2*^-/-^*Zbtb25*^-/-^ cells in 2iLIF. **(C)** Diagram of ZBTB2 constructs used for Y2H analysis; + and – indicates positive and negative interactions as in (**E**,**F**). **(D-G)** Colony growth of strains expressing indicated full length and deletion proteins. Bait constructs are vertical (**D**,**E**) and horizontal (**F**,**G**), and prey constructs are horizontal (**D**,**E**) and vertical (**F**,**G**). **(H**,**I)** Volcano plot of protein enrichments in AP-MS of ZBTB2-AVI in *WT* compared to *Znf639*^-/-^cells in 2iLIF. Color code as in Fig. 2A (**H**). Bait-normalized interaction changes in WT compared to *Znf639* mutant cells (**I**). * indicates p-value<0.01 and ** p-value<0.001.

We, therefore, turned to yeast-two-hybrid (Y2H) assays. First, we addressed dimerizing properties of ZBTB2. We detected homodimerization of ZBTB2’s btb domain and heterodimerization with ZBTB25 (Fig. 2C, D). To our surprise, we also found a strong interaction of ZBTB2’s linker region with ZBTB2’s btb domain but not with the full-length ZBTB2 protein (Fig. 2C, D). The linker is 136 amino acid in length and predicted to be unstructured. We generated linker region deletions and found that the segment immediately adjacent to the btb domain (link-D1) mediates interaction with the btb domain (Fig.2C, E), suggesting the existence of an extended btb domain structure. We therefore used this extended btb domain (btb-link) in further assays. Next, we tested the direct interaction of ZBTB2 with the HiRA complex subunits HIRA, CABIN1, and UBN2. We found that UBN2, but not HIRA or CABIN1, binds full-length ZBTB2 and the btb-link domain, but neither the isolated btb domain or linker region (Fig. S2C). Deletion analysis of UBN2 identified a minimal interacting region of 68 amino acids that is outside of annotated domains (Fig. S2D, E). The btb-link domain therefore mediates homo- and heterodimerization with ZBTB2 and ZBTB25, respectively, and interaction with UBN2.

We then tested binding to the NuRD subunits RBBP4, MBD3, MTA2, MTA3, HDAC1, GATAD2A, and GATAD2B. Based on our AP-MS results (Fig. 2B), we expected that Znf1 would mediate such an interaction, yet the only direct interactions we identified were between GATAD2A and ZBTB2, and between GATAD2B and both, ZBTB2 and the btb-link domain (Fig. S2F, G). No interactions were found with Znf1, the btb domain or the linker region. Additional Y2H assays revealed that the C-terminal half of GATAD2B, which includes its Gata-type Znfs, mediates interaction with the btb-link domain (Fig. S2H, I).

As the direct interactions with both UBN2 and GATAD2B were mediated by the btb-link domain, but not the btb domain or linker alone, we sought to determine the minimal btb domain extension required for either interaction. Through serial truncations we found that a 44 amino acid extension was necessary for both (Fig. 2C, F). This, together with the direct interaction between the conserved btb domain and this 44 amino acid fragment (Fig. 2C, E) strongly suggests a functionally essential structural extension of ZBTB2’s btb domain.

Further Y2H assays identified a direct interaction between ZNF639 and ZBTB2’s Znf1 domain (Fig. 2G), as expected by the AP-MS data (Fig. 2B). Znf1 was required for the interaction with NuRD in AP-MS (Fig. 2B), but did not mediate any direct interaction with NuRD subunits (Fig. S2F, G). Therefore, we wondered whether ZNF639 is required to stabilize or enhance the interaction between NuRD and ZBTB2, which is mediated by the extended btb domain. To test this hypothesis, we performed AP-MS of ZBTB2-AVI in *wt* and *Znf639*^-/-^ mESCs (Fig. S2J, Table S2), and found a ∼8-fold reduction in NuRD interaction upon loss of *Znf639* (Fig. 2H, I).

In summary, our AP-MS and Y2H experiments show that the btb domain of ZBTB2 mediates homodimerization and heterodimerization with ZBTB25, that an extended btb domain recruits the UBN2-HiRA complex through the UBN2 subunit, and that the interaction with MBD3-NuRD is direct via btb-link binding GATAD2A/B, but also requires Znf1 recruiting ZNF639.

### Recruitment of HiRA is a unique property of ZBTB2, while GATAD2A/B interaction is a conserved feature of TF-associated btb domains

Interaction of a btb-TF with the HiRA complex has not been reported before, while the interaction with NuRD has been previously shown for ZBTB7A (Masuda et al., 2016). We therefore wondered if HiRA and NuRD recruitment would be conserved across btb-TFs. We first attempted to identify additional HiRA-interacting btb domain proteins by performing UBN2 and UBN1 AP-MS in mESCs. UBN2 pull-down identified the HiRA components CABIN1, HIRA, and UBN1. The only TFs we recovered were ZBTB2 and its partner ZNF639 (Fig. 3A), but not ZBTB25. In the UBN1 AP-MS we detected ZBTB2 and MEF2D (Fig. S3A), a known direct interactor of CABIN1 (Youn et al., 1999).

**Figure 3:**
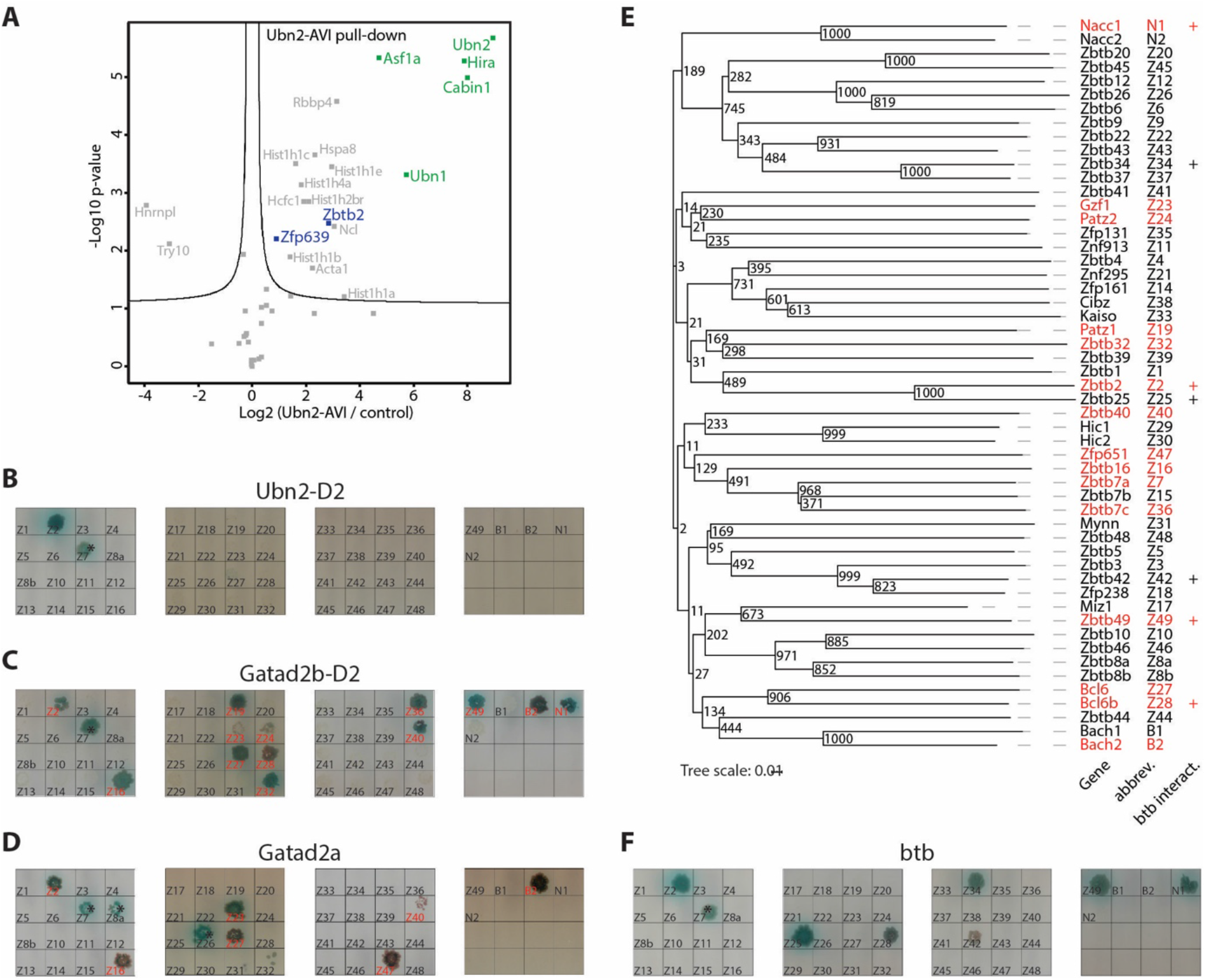
Recruitment of HiRA is a unique property of ZBTB2, while GATAD2A/B interaction is a conserved feature of TF-associated btb domains. **(A)** Volcano plot of protein enrichments in AP-MS of UBN2-AVI compared to control BirA-expressing cells in 2iLIF. ZBTB2 and partner TFs are indicated in blue and HiRA subunits in green. (**B-D**,**F**) Colony growth of strains expressing extended btb domains of *Zbtb* (Z#), *Bach* (B#) and *Nacc* (N#) bait constructs, and Ubn2-D2 **(B)**, Gatad2b-D2 **(C)**, Gatad2a **(D)** and ZBTB2 btb domain (**F**) prey constructs. GATAD2A/B interactors are indicated in red. Asterisks mark autoactivating bait constructs. **(E)** Phylogenetic tree of TF-associated btb domains with bootstrap values. GATAD2A/B interactors, based on Fig. 2C,D and (Masuda et al., 2016) are indicated in red. + indicates btb domains dimerizing with ZBTB2.

As not all btb-TFs are expressed in mESCs, we turned to Y2H to systematically assay ability to bind to UBN2 or GATAD2A/B. We performed Y2H screens of the btb domains of the 49 *Zbtb* factors, *Nacc1/2*, and *Bach1/2*, extending the conserved btb domains by at least 60 amino acids, in case other btb domains would possess extended structures similar to *Zbtb2*. Strikingly, only ZBTB2’s btb-link domain interacted with UBN2 (Fig. 3B, Fig. S3B), corroborating the UBN1/2 AP-MS data to support that HiRA recruitment by ZBTB2 is unique. Surprisingly, we identified btb domains of 14 btb-TFs to interact with GATAD2A or GATAD2B (Fig. 3C, D, Fig. S3B, C). We were not able to confirm binding to ZBTB7A’s btb domain (Masuda et al., 2016) due to autoactivation in the Y2H assays (Fig. S3B, C). Taken together, at least 15 out of 54 btb-TFs bind to NuRD subunits, suggesting that GATAD2A/B interaction is a common function of TF-associated btb domains.

We wondered whether the GATAD2A/B interacting domains would be phylogenetically related. Phylogenies based on the complete sequence of *Zbtb* TFs do not reflect the similarities within the btb domains (Siggs and Beutler, 2012) because of the influence of the Znf domains. We therefore constructed a phylogenetic tree based on btb domain sequences (Fig. 3E, Table S6). Although confidence for the evolutionary older branches was low, GATAD2A/B -interacting btb domains do not form a clade, but are scattered throughout the tree, including the *Bach* and *Nacc* clades. This shows that GATAD2A/B recruitment is an ancestral property of TF-associated btb domains.

To test for heterodimerization with ZBTB2 and if this correlates with binding to GATAD2A/B, we repeated the Y2H btb domain family screen using ZBTB2’s btb domain as bait (Fig. 3F, Fig. S3B). This identified 6 btb domains heterodimerizing with ZBTB2, of which 3 also bound to GATAD2A/B. Specificity for heterodimerization and GATAD2A/B binding are therefore separate properties of btb domain.

Although ZBTB25’s btb domain heterodimerizes with ZBTB2, it did not bind UBN2 or GATAD2A/B (Fig. 3B-F). We therefore hypothesized that binding to ZBTB25 is not relevant to ZBTB2’s role in cell fate transition. To test this, we characterized the phenotype of *Zbtb25*^-/-^ and *Zbtb2*^-/-^ *Zbtb25*^-/-^ mESCs (Fig. S3D) in the 2iLIF to Serum-LIF transition and compared it to *Zbtb2*^-/-^ cells. In fact, loss of *Zbtb25* did not delay Nanog>GFP downregulation or modify the phenotype of *Zbtb2* single mutants (Fig. S3E, F).

In summary, we tested the conservation of UBN2 and GATAD2A/B interaction by AP-MS and Y2H. While UBN2 recruitment appears to be unique to ZBTB2, GATAD2A/B interaction is a common and ancestral feature of btb-TFs.

### ZBTB2 interacts with the Ep400 complex in a HiRA-independent manner

While evaluating the functionality of tagged *Zbtb2* constructs for AP-MS, we found that overexpression of ZBTB2 fused with an extended C-terminal 3xHA-AVI-3xFLAG-tag (HAF-tag) caused a delay in *Nanog>GFP* reporter downregulation (Fig. 4A). This phenotype is opposite to ZBTB2-AVI overexpression but similar to loss of *Zbtb2*, suggesting that ZBTB2-HAF acts dominant negative. To understand the underlying molecular defect, we compared the interactomes of the ZBTB2-AVI and ZBTB2-HAF fusion proteins. To our surprise, we found that the entire Ep400 complex copurified with the dominant negative ZBTB2-HAF, while none of the other interactors was lost (Fig. 4B). This prompted us to look more carefully at the ZBTB2-AVI AP-MS data and consistently found Ep400 subunit peptides across independent experiments (Table S5), suggesting that ZBTB2-HAF stabilizes a physiological, but transient or weak interaction.

**Figure 4:**
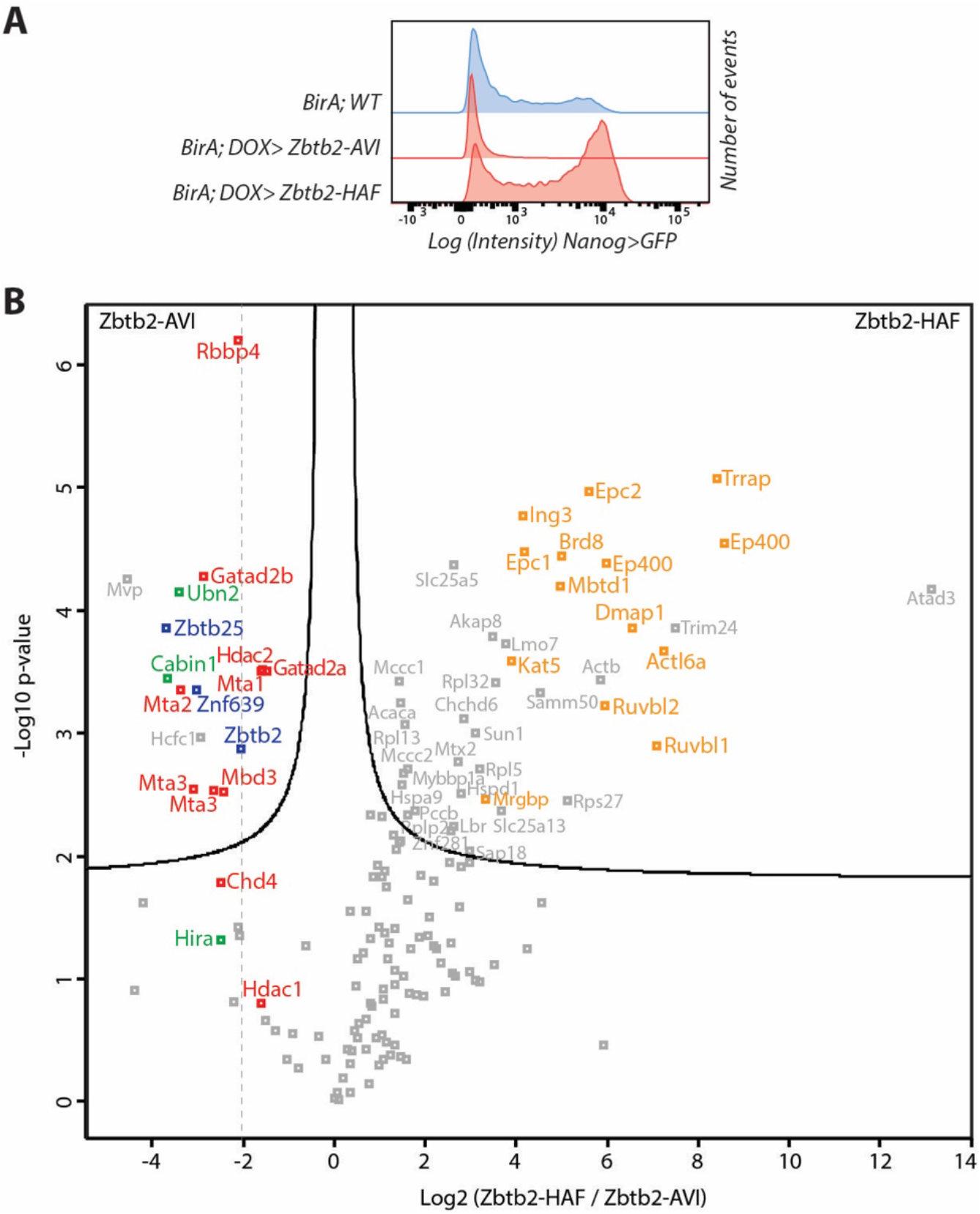
A dominant negative *Zbtb2* construct stabilizes the interaction with the Ep400 complex. **(A)** *Nanog>GFP* intensities upon Zbtb2-AVI or Zbtb2-HAF induction for 3 days in N2B27 +CHIR+bFGF. **(B)** Volcano plot of protein enrichments in AP-MS of ZBTB2-HAF compared to ZBTB2-AVI. ZBTB2 and partner TFs are indicated in blue, NuRD subunits in red, HiRA subunits in green, and Ep400 subunits in orange.

Ep400 incorporates H2A.Z/H3.3 histones into chromatin (Pradhan et al., 2016). We therefore hypothesized that association of Ep400 with ZBTB2 could be mediated by the HiRA complex, which is an H3.3 chaperone (Tagami et al., 2004). To test this, we performed AP-MS of ZBTB2-HAF in *wt* and in *Ubn2*^-/-^ mESCs (Fig. S4A). As expected, lack of UBN2 caused loss of the HiRA interaction, but the association with Ep400 was not affected (Fig. S4B). Interestingly, we noted a substantial increase of ZBTB25 in ZBTB2 AP-MS in *Ubn2*^-/-^ cells (Fig. S4B). UBN2 and ZBTB25 may therefore compete for interaction with ZBTB2. Together with the inability of ZBTB25 to interact with UBN2 (Fig. 3B), this supports the idea that ZBTB25 is a negative regulator of HiRA recruitment by ZBTB2. We therefore conclude that ZBTB2 interacts weakly or transiently with Ep400 and independent of HiRA co-binding.

### Ep400 and Znf639/NuRD/HiRA constitute independent Zbtb2 functional modules

To address the functional role of ZBTB2’s protein interactions, we set out to compare the loss of function phenotype of *Zbtb2* with that of its binding partners. Depletion of the Ep400 complex subunits *Ep400* and *Kat5* causes loss of mESC self-renewal (Fazzio et al., 2008), while knockout of *Mbd3* (Kaji et al., 2006) or *Hira* (Leeb et al., 2014) induces resistance to exit from the mESC state during differentiation. However, these phenotypes may arise from pleiotropic, *Zbtb2*-unrelated roles. As the ZBTB2-HiRA interaction is dependent on UBN2, and the ZBTB2-NuRD interaction is stabilized by ZNF639, we reasoned that analysis of *Ubn2*^-/-^ and *Znf639*^-/-^ cells might reveal *Zbtb2*-specific functions. Compared to *Zbtb2* mutants, *Znf639*^-/-^ and *Ubn2*^-/-^ mESCs showed moderate delays in *Nanog>GFP* downregulation upon Serum-LIF transition (Fig. 5A, B). Although weak, these phenotypes are consistent with *Znf639* and *Ubn2*, and by extension NuRD and HiRA, cooperating with *Zbtb2* in cell fate transitions.

**Figure 5:**
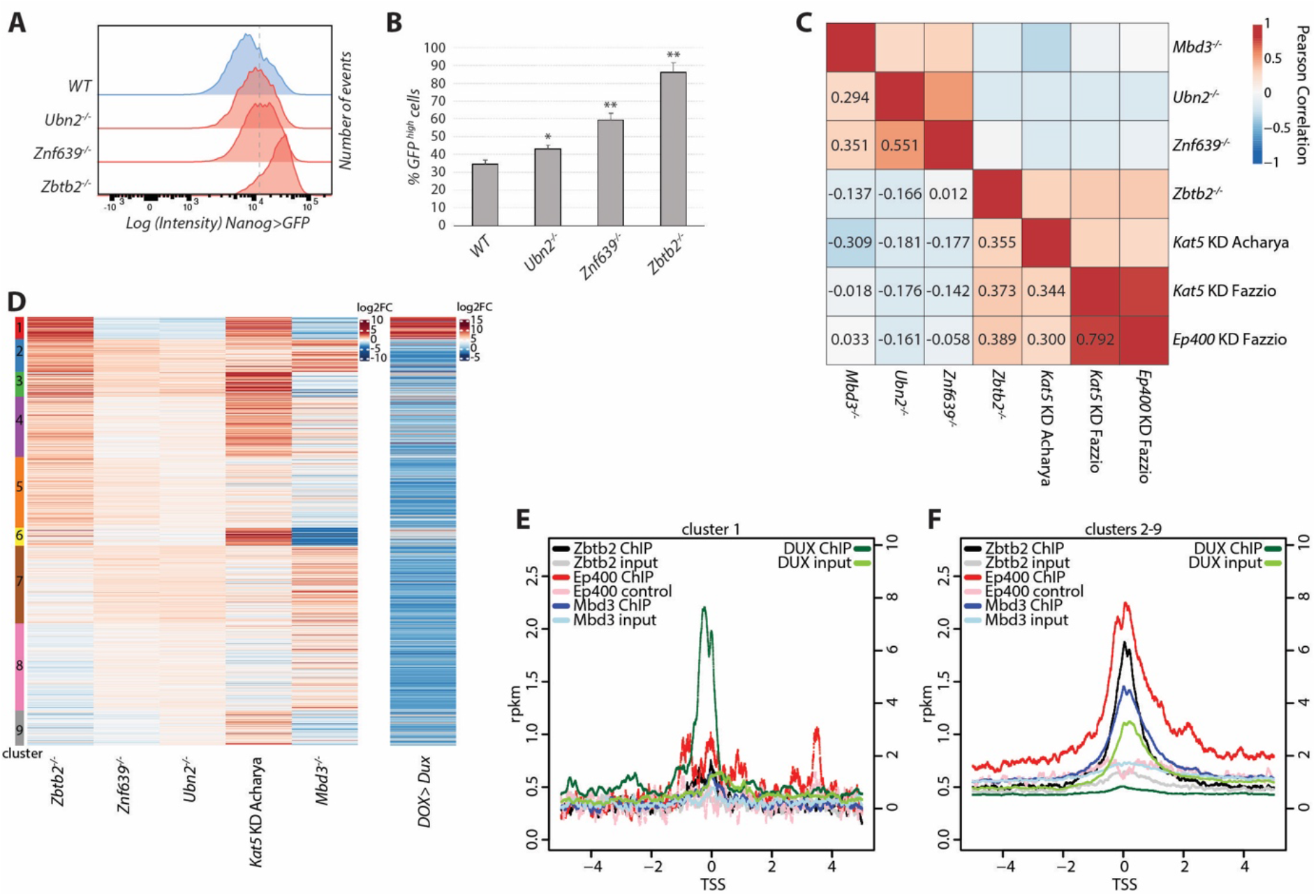
Ep400 and ZNF639/NuRD/HiRA constitute independent functional modules. **(A**,**B)** *Nanog>GFP* intensities 3 days in Serum-LIF of indicated genotypes (**A**). Dashed line indicates the threshold for quantification of GFP-high cells as average and SD of biological triplicates (**B**). * indciates p-values<0.01 and ** p-values<0.001 compared to the *WT* control. **(C)** Pairwise Pearson correlations of differential gene expression relative to respective control cells after 48h in Serum-LIF for *Ubn2*^-/-^, *Znf639*^-/-^ and *Zbtb2*^-/-^ mutant cells, and for *Mbd3*^-/-^ (Reynolds et al., 2012), *Kat5* KD (Acharya et al., 2017; Fazzio et al., 2008) and *Ep400* KD (Fazzio et al., 2008) in Serum-LIF. Only genes deregulated in *Zbtb2*^-/-^ cells were considered (1420 genes). **(D)** k-means clustering of differential gene expression as in Fig. 2C and upon *Dux* overexpression (Hendrickson et al., 2017). **(E**,**F)** ZBTB2 (Karemaker and Vermeulen, 2018), EP400 (Chen et al., 2015) and MBD3 (Bornelöv et al., 2018) (left scale), and DUX (Hendrickson et al., 2017) (right scale) ChIP-seq reads per kilobase of transcript per million mapped reads (rpkm) of centered on transcriptional start sites (TSSs) of cluster 1 (**E**) and cluster 2-9 genes (**F**).

To investigate these functional relationships further, we performed RNAseq of *Zbtb2*^-/-^, *Znf639*^-/-^ and *Ubn2*^-/-^ mESCs after 48h in Serum-LIF. Consistent with *Nanog* reporter phenotypes, pre-implantation epiblast-enriched transcripts were upregulated and post-implantation epiblast-specific genes were downregulated in *Znf639* and *Ubn2* mutant cells in Serum-LIF, although to a lesser extent than observed in *Zbtb2* knockout cells (Fig. S5A). To describe *Zbtb2*’s transcriptional role, we focused on genes significantly changing upon loss of *Zbtb2* and included published transcriptome data of *Mbd3*^-/-^ (Reynolds et al., 2012), *Ep400* and *Kat5* knock-down (KD) (Acharya et al., 2017; Fazzio et al., 2008) mESCs in Serum-LIF (Table S3). We found that transcriptional alterations in *Zbtb2*^-/-^ cells correlated with those upon knockdown of *Ep400* (R=0.39) and *Kat5* (R=0.37 and 0.36), showing a mechanistic relationship between *Zbtb2* and the Ep400 complex (Fig. 5C). Changes in *Mbd3* mutants, in contrast, correlated with those in *Znf639* knockout cells (R=0.36), corroborating the strong reduction of NuRD binding to ZBTB2 in *Znf639* mutants. Furthermore, alterations in *Mbd3, Znf639*, and *Ubn2* mutants correlated reciprocally, pointing towards a functional ZNF639/NuRD/HiRA unit (Fig. 5C). Strikingly, there was no general correlation between *Zbtb2*/*Ep400*-dependent and *Znf639*/*Mbd3*/*Ubn2*-dependent gene expression, raising the hypothesis that Ep400 and ZNF639/NuRD/HiRA cooperate with ZBTB*2* within functionally independent modules.

For more detailed insight, we performed k-means clustering of differential gene expression (Fig. 5D). This identified cluster 1 genes to be upregulated in *Zbtb2*^-/-^ and *Kat5* KD cells and downregulated in *Znf639*^-/-^, *Mbd3*^-/-^, and *Ubn2*^-/-^ cells. Cluster 1 contains the strongest changing genes in *Zbtb2*^-/-^ cells and closer inspection revealed them to be enriched for 2C-like genes, as corroborated by the correlation with DUX-induced genes (Fig. 5D) (Hendrickson et al., 2017). Such opposite functions are consistent with independent complexes competing for limiting ZBTB2 amounts, where loss of the ZNF639/NuRD/HiRA complex would lead to an increase in Ep400/ZBTB2 dependent 2C-like gene repression.

Taken together, our findings show that *Zbtb2, Znf639* and *Ubn2* loss of function cause a developmental delay that is reflected in the deregulation of pre- and post-implantation epiblast specific genes. However, a systematic analysis of ZBTB2 target genes revealed that gene expression changes upon loss of *Ep400* and *Kat5* correlate best with *Zbtb2* mutants, while alterations upon loss of *Znf639, Ubn2*, and *Mbd3* correlate reciprocally, but not with *Zbtb2* mutants. This suggests the existence of two functionally distinguishable ZBTB2 modules, one associated with the Ep400 complex, causing most *Zbtb2*-dependent gene expression changes, and the other with ZNF639/NuRD/HiRA. Opposite roles in regulating 2C-like genes expression suggest that these modules act antagonistically within one protein complex, or independently in competing biochemical complexes.

### ZBTB2 inhibits 2C-like gene transcription indirectly but binds and represses its own promoter

The Ep400 complex represses expression of 2C-like genes without binding to their regulatory DNA sequences (Rodriguez-Terrones et al., 2018). We therefore wondered if ZBTB2 is similarly depleted at 2C-like genes. As ZBTB2 mostly binds to promoters (Karemaker and Vermeulen, 2018), we analyzed ZBTB2, EP400, KAT5 (Chen et al., 2015), MBD3, and CHD4 (Bornelöv et al., 2018) occupancy at promoters of genes belonging to cluster 1 (2C-like genes) or clusters 2-9 (Fig. 5E, F, and Fig. S5B, C). We found that these factors are depleted at cluster 1 promoters, as opposed to DUX which is highly enriched, as expected (Hendrickson et al., 2017). Therefore, 2C-like gene repression by ZBTB2/Ep400 is either indirect through the regulation of other genes, or mediated through DNA-binding independent mechanisms.

Nevertheless, ZBTB2 is enriched at promoters, where it colocalizes with EP400, KAT5, MBD3, and CHD4 (Fig. S5D). Therefore, we wondered what the activity of ZBTB2 on bound genes is. One of the most prominent ZBTB2 ChIP-seq peaks lays on the promoter of *Zbtb2* itself (Fig. S5E). As some of our *Zbtb2* mutants generated *Zbtb2* transcripts that were detectable by quantitative polymerase chain reaction (qPCR) (Fig. S1C, Fig. S5E), this gave us the opportunity to test the activity of ZBTB2 on its own promoter. Mutation of *Zbtb2* leads to increased *Zbtb2* transcript levels and overexpression of a *Zbtb2* construct not detected by our qPCR probes leads to decreased endogenous *Zbtb2* transcript levels (Fig. S5F). In conclusion, ZBTB2 represses transcription either via promoter binding, as is the case for autoregulation, or via indirect mechanisms, as is the case for 2C-like genes.

## DISCUSSION

In this study, we exploited a genetic screen in mESCs to mechanistically and functionally characterize the btb-TF ZBTB2, gaining insights that are broadly applicable to btb-TFs.

We made use of a sensitized set-up and took advantage of a loss-of-function and gain-of-function haploid mESC library, identifying previously described and novel regulators of exit from pluripotency (Fig. 1A). We validated the role of *Zfp42*/*Rlim, Nmt1*, and *Zbtb2* in an independent cell line and using an unrelated reporter (Fig. 1B, C, D), confirming their role in mESC differentiation. Previous studies of *Zfp42* (Masui et al., 2008; Scotland et al., 2009) led to the prevailing idea that *Zfp42* is dispensable for pluripotency or development, to the point that *Zfp42*^-/-^ cells have been used in genetic screens for the exit from pluripotency (Leeb et al., 2014; Villegas et al., 2019; Yang et al., 2012). Our data also supports the relationship between *Rlim* and *Zfp42* (Gontan et al., 2012) and calls for further investigation of the function of this genetic axis in pluripotency. *Nmt1* is the major N-Myristoyltransferase in mESCs and is essential in early mouse development (Yang et al., 2005). As some of its substrates are known (Thinon et al., 2014), we tested two possible modes of action, via FRS2 (Kouhara et al., 1997) and MAPK-signaling, or via LAMTOR1 and TFE3 localization (Villegas et al., 2019). While we were not able to detect changes in ERK phosphorylation (Fig. S1E), we noticed a clear increase in nuclear TFE3 (Fig. 1E), which is compatible with a TFE3-dependent differentiation delay.

With the aim of gaining fundamental insights into transcriptional regulation of cell state transition, we focused our work on the btb-TF *Zbtb2*. A previous report showed morphological delay upon LIF removal in *Zbtb2*^-/-^ cells (Karemaker and Vermeulen, 2018), but did not characterize this phenotype further. We found that *Zbtb2*^-/-^ naïve mESCs are defective in differentiating into the EpiLC and Serum-LIF cell states (Fig. 1F). Upon transitioning to Serum-LIF, in particular, *Zbtb2*^-/-^ cells fail to timely upregulate post-implantation epiblast genes and to down-regulate pre-implantation epiblast genes (Fig. 1G). To understand the underlying molecular mechanism, we performed a thorough biochemical analysis of ZBTB2-containing protein complexes and showed (1) that ZBTB2’s btb domain mediates homodimerization, heterodimerizaton with ZBTB25, interaction with UBN2-HiRA via UBN2, and interaction with MBD3-NuRD via GATAD2A/B, (2) that ZBTB2 first Znf interacts with ZNF639 and that this interaction is required to establish or stabilize binding of ZBTB2 to NuRD, and (3) that ZBTB2 interacts weakly or transiently with the Ep400 complex. Ep400 incorporates H3.3 containing histones into chromatin (Pradhan et al., 2016), for which HiRA is a chaperone (Tagami et al., 2004). Nevertheless, we show that Ep400 recruitment to ZBTB2 is independent of HiRA (Fig. S4B). Although we identified binding to Ep400 using a dominant negative ZBTB2 construct (ZBTB2-HAF), we propose that this is a physiologically relevant interaction, because: (1) low levels of Ep400 subunits are detectable in AP-MS of the functional ZBTB2-AVI construct (Table S5), (2) transcriptional changes upon loss of *Zbtb2* and *Ep400*/*Kat5* correlate (Fig. 5C, D), (3) a direct interaction between ZBTB2 and the Ep400 subunit KAT5 was identified by high-throughput Y2H screening (Luck et al., 2020).

These molecular insights have important implications for our understanding of histone variant deposition. The prevailing models for HiRA recruitment hypothesize a transcription-coupled mechanism (Sarai et al., 2013) or intrinsic affinity for nucleosome-free DNA (Ray-Gallet et al., 2011). As ZBTB2 is able to recruit HiRA and to localize at promoters, we propose that HiRA recruitment at transcription start sites can be mediated by ZBTB2. The HiRA complex binds H3.3 (Tagami et al., 2004), yet there is no known HiRA-associated chromatin remodeler for H3.3 deposition, such as ATRX for the H3.3 chaperone DAXX (Grover et al., 2018). Our work provides the first physical link between HiRA and a chromatin remodeler. The hypothesis that HiRA might be coupled to Ep400 is further supported by Ep400’s preference for H2A.Z/H3.3 nucleosome deposition (Pradhan et al., 2016).

While btb domain-containing TFs are acknowledged to play biologically important and disease-relevant roles (Chevrier and Corcoran, 2014), a general understanding of their evolution and molecular function is lacking. Building on our characterization of ZBTB2’s interactome, we systematically assayed the biochemical properties of btb-TFs. Using Y2H we found that UBN2 recruitment is a unique feature of ZBTB2. GATAD2A/B binding, instead, is shared by at least 15 btb-TFs and is therefore the most common feature of btb-TFs reported to date. Instances of this interaction are present in the *Nacc* family, in the *Bach* family, and in many apparently unrelated branches of the *Zbtb* family (Fig. 3E), suggesting that GATAD2A/B binding is an ancestral property of btb-TFs. Intriguingly, the Human Reference Protein Interactome Mapping Project (Luck et al., 2020) identified the btb-TFs ZBTB1, ZBTB2, ZBTB8A, ZBTB14, and BACH2 to directly interact with KAT5, suggesting that Ep400 recruitment might also be a conserved property of btb-TFs. The dimerization specificity of btb domains is incompletely understood and has been proposed to be enforced by a quality control mechanism (Mena et al., 2018). In contrast, our Y2H data for ZBTB2 heterodimerization shows that dimerization specificity is a btb domain-intrinsic property and unrelated to interaction with other partners. For example, ZBTB25’s btb domain heterodimerizes with ZBTB2’s btb domain (Fig. 3F), but does not interact with GATAD2A/B or UBN2, while ZBTB2’s btb domain does so (Fig. 3B-D). This raised the possibility that ZBTB25 might modulate ZBTB2’s avidity for HiRA and NuRD, which was confirmed, in the case of HiRA, by the competition between UBN2 and ZBTB25 for ZBTB2 binding (Fig. S4B). Nevertheless, this regulatory mechanism does not seem to play an important role, as the lack of *Zbtb25* does not affect differentiation in either *WT* or *Zbtb2*^-/-^ cells (Fig. S3E, F).

The linker region between the btb domain and the first Znf of ZBTB proteins is not conserved (Stogios et al., 2005) and, although it can mediate protein interactions, is usually considered to work as a flexible unstructured linker (Maeda, 2016). We found that the first 44 amino acids of ZBTB2’s linker interact with the conserved btb domain, but not with the extended btb domain (Fig. 2E), and that extension by these 44 amino acids is necessary for the btb domain to bind UBN2 and GATAD2B (Fig. 2F). We interpret this as evidence for a structured extension of the btb domain of ZBTB2. As ZBTB7A does not require a btb domain extension to interact with GATAD2B (Masuda et al., 2016), we hypothesize that this is a unique feature of ZBTB2 that evolved with its ability to bind UBN2.

To assess the function of the characterized ZBTB2 interactions, we generated *Znf639*^-/-^ and *Ubn2*^-/-^ mESCs and analyzed their differentiation phenotypes and transcriptional alterations. *Znf639*^-/-^ and *Ubn2*^-/-^ mESCs show a delay in *Nanog>GFP* downregulation (Fig. 5A, B) that, although weaker than in *Zbtb2* mutants, is consistent with the reported phenotypes of *Mbd3*^-/-^ (Kaji et al., 2006) and Hira^-/-^ cells (Leeb et al., 2014). At the transcriptional level, *Mbd3, Ubn2* and *Znf639* mutants correlate reciprocally (Fig. 5C), behaving as a functional unit, which is consistent with the role of ZNF639 in stabilizing NuRD interaction (Fig. 2H). Although we do not know whether ZNF639, UBN2 and NuRD simultaneously bind ZBTB2, this suggests that they work synergistically, rather than regulating independent genes. *Znf639*/NuRD/HiRA module mutants show no transcriptional correlation with *Zbtb2*^-/-^ cells, suggesting that ZBTB2 directs gene regulation predominantly through another interactor, such as the Ep400 complex. In fact, *Zbtb2*-specific expression changes correlate with those upon *Ep400* and *Kat5* depletion (Fig. 5C), pointing to the existence of two separate ZBTB2 effector modules, one associated with Ep400, and the other with ZNF639/NuRD/HiRA. The phenotypic convergence of *Zbtb2, Znf639, Mbd3* and *Ubn2*/*Hira* on promoting exit from naïve pluripotency might be due to coherent regulation of pre- and post-implantation specific genes (Fig. S5A). The role of Ep400 in driving mESC differentiation remains to be determined, since *Ep400* and *Kat5* are essential for ESC self-renewal (Fazzio et al., 2008).

ZBTB2 and the Ep400 complex (Rodriguez-Terrones et al., 2018) repress 2C-like genes, while ZNF639, MBD3, and UBN2 promote their transcription (Fig. 5D). Nevertheless, all these factors are depleted at the promoters of 2C-like genes (Fig. 5E, F, Fig. S5B, C), demonstrating that this regulation is indirect. The underlying mechanism remains to be determined. Since the H3.3 histone chaperone DAXX/ATRX directly inhibits 2C-like gene expression (Elsässer et al., 2015; Sadic et al., 2015), this mechanism may involve ZBTB2 modulating H3.3 dynamics through Ep400 and HiRA. A similar mechanism may contribute to the regulation of 2C-like genes by the canonical H3 chaperone CAF1 (Ishiuchi et al., 2015). However, ZBTB2 can also repress transcription in a sequence-specific and direct manner, which we demonstrate by taking advantage of a prominent ZBTB2 ChIP-seq peak at the *Zbtb2* promoter and showing that ZBTB2 regulates its own transcription in a negative feedback-loop (Fig. S5F).

In summary, this study presents a detailed biochemical and transcriptional analysis of ZBTB2, and identifies how chromatin modifiers and histone chaperones are recruited by this TF during cell state transition. We use these molecular insights to systematically analyze btb-TFs and to propose a comprehensive concept for their evolution and function. This work will serve as a resource for the study of btb-TFs and inspire further systematic approaches to this important family of TFs.

## ACKNOWLEDGMENTS

We thank Hans-Rudolf Hotz (FMI) for help with the phylogenetic analysis; H. Kohler (FMI) for cell sorting; V. Iesmantavicius (FMI) for help with computational analysis of mass spectrometry data; Min Jia and Jeff Chao (FMI) for providing LIF; Austin Smith (University of Exeter) for providing TNG-A cells. D.O. was supported by EMBO (ALTF 1632-2014) and Marie Curie Actions (LTFCOFUND2013, GA-2013-609409). Research in the lab of J.B. is supported by the Novartis Research Foundation.

## AUTHOR CONTRIBUTIONS

D.O. and J.B. conceived the study; D.O. designed, performed, and analyzed experiments; S.P. performed Y2H experiments; D.H. mass spectrometry; U.E. haploid mESCs generation and mutagenesis; A.F.B. analyzed the screen data; S.A.S. supervised sequencing library generation and sequencing; J.B. performed computational analysis; D.O. wrote the paper.

## MATERIALS AND METHODS

### Genetic screen

175*10^6^ haploid mESCs B1A4 Oct4>GFP-Puro (Elling et al., 2011) were transduced with each of the retroviruses reFlipROSAβgeo(Cre)*0, reFlipROSAβgeo(Cre)*+1, reFlipROSAβgeo(Cre)*+2, rsFlipROSAßgeo*+2 (Schnütgen et al., 2008), selected with Neomycin, and frozen. After thawing, cells were allowed to recover in Serum-LIF (GMEM (Sigma), 10% fetal bovine serum (Sigma), 1mM sodium pyruvate (Gibco), 2mM L-glutamine (Gibco), 0.1mM non-essential amino acids (Gibco), 0.1mM 2-mercaptoethanol (Sigma), and 1000U/mL mLIF (Chao lab, Basel)) for 24 hours and then transferred on day 0 to N2B27 (DMEM/F12 medium (Life Technologies) and Neurobasal medium (Gibco) 1:1, N2 supplement 1/200 (homemade), B-27 Serum-Free Supplement 1/100 (Gibco), 2mM L-glutamine (Gibco), 0.1mM 2-mercaptoethanol (Sigma)) + 3µM CHIR99021 (Steward lab, Dresden). Fresh medium was provided every 2 days and the cells were replated every 4 days until day 13, when the medium was changed to N2B27 + CHIR + Puromycin. Half of the cells were harvested on day 16, then again on day 20, and finally all on day 23. Cell pellets were digested overnight at 56°C in 10mM TrisHCl pH7.5, 10mM EDTA, 10mM NaCl, 0.5%SDS, 1mg/mL proteinase K (Macherey-Nagel) and then with 0.1 mg/mL RNaseA (Qiagen) for 2h at 37°C. DNA was ethanol precipitated, washed twice with ethanol 70%, and resuspended in H_2_O. 5ug of DNA were digested with DpnII or MseI (NEB) for 5h at 37°C, purified with the PCR cleanup kit (Qiagen), ligated in 250μL with 3μL of T4 ligase (NEB) at 16°C for 36h, and purified with the PCR cleanup kit (Qiagen). Half of the eluate was redigested with NheI and half with PvuII (NEB), the reactions were pooled and purified by PCR cleanup kit (Qiagen) eluted in 50μL. 10μL of eluate were used as a template for PCR with KOD (Takara) with extension time 1’10’’, annealing temperature 58°C, for 35 cycles, with primers aatgatacggcgaccaccgagatctacacGCCAGTCCTCCGATTGA and caagcagaagacggcatacgagatBBBBBBAGTTCCTATTCCGAAGTTCCTATTCTCTA (B=barcode). PCRs were purified with the PCR cleanup kit (Qiagen), and subjected to next generation sequencing with sequencing primer TGATTGACTACCCGTCAGCGGGGGTCTTTCA and indexing primer TATACTTTCTAG+A+GAATAGGAACTTCGGAATA+G+GAACT (+N = LNA modification).

Sequence reads were processed to remove adaptors (Msel TTAA and Dpnll GATC) and then mapped to the mouse reference genome (mm9 only chromosomes 1 to 19, X, Y and M) using Bowtie version 1.0.0 with parameters -v 3 -m 1 —best —strata. Output SAM files were sorted and converted to BAM files using samtools version 0.1.19-44428cd. Genomic tracks were generated using *genomeCoverageBed* from bedtools version 2.25.0 using only the first base pair position of each read according to the strand. The number of insertions per gene (using only insertions supported by more than one read) normalized to 10,000 reads per library was generated using the ENSEMBL gtf annotation Mus_musculus.NCBIM37.67. Most of this analysis was processed using the unix command *awk*. The R version 3.3.2 was used to compute a mean log2 fold change (using a pseudocount of 0.5) contrasting each group to the corresponding control and a z-score within each library (Table S1).

### Cell culture

TNG-A mESCs (Chambers et al., 2007) were cultured in 2iLIF (N2B27, 1µM PD0325901 (Steward lab, Dresden), 3µM CHIR99021, and 1000U/mL mLIF) on gelatin-coated tissue culture plates. For medium switch experiments, cells were washed with PBS, detached with Accutase (Sigma), centrifuged for 3’ at 300g in DMEM/F12-0.1% BSA, resuspended in the new medium, counted, and replated. In experiments with doxycycline inducible constructs, 1μg/mL doxycycline (Sigma) was added to the final medium. Cells were transitioned to N2B27 + 3µM CHIR99021 + 12 ng/ml bFGF (Smith lab, Cambridge) on gelatin coated plates at a density of 2’500/cm^2^, or to EpiLC medium ((N2B27 base, 20 ng/ml activin A, 12 ng/ml bFGF (Smith lab, Cambridge), and 1% KSR (Life Technologies)) on fibronectin coated plates at a density of 25’000/cm^2^, or to Serum-LIF on gelatin coated plates at a density of 2’500/cm^2^. At the moment of analysis, cells were washed with PBS, resuspended with trypsin (Life Technologies), centrifuged for 3’ at 300g in DMEM/F12-0.1% BSA, resuspended in DMEM/F12-0.1% BSA, and flowed on an LSRII SORP Analyzer (Becton Dickinson). Percentage of GFP-high cells was quantified with BD FACSDiva 8.0.1 and flow profiles were made with FlowJo (FlowJo, LLC).

### Mutant cell lines and overexpression constructs

A TNG-A clone stably expressing Cas9 (TbC1) was derived by transfection of PB-LR5.1-EF1a-bsdr2ACas9 (derived from pPB-LR5.1-EF1a-puro2ACas9, gift of Kosuke Yusa, Wellcome Trust Sanger Institute) and PBase (PiggyBac Transposase, (Betschinger et al., 2013)). To generate KO cell lines, TbC1 cells were reverse transfected with 400ng of U6>sgRNA plasmids (George Church, Addgene plasmid #41824) according to Table S2 and 3μL Lipofectamin 2000 (Life Technologies). Single cells were sorted in 96-well plates in 2iLIF and expanded. For constitutive or inducible expression, the cDNA of the gene of interest was cloned in pPB-CAG-DEST-pgk-hph (CAG>) (Betschinger et al., 2013) or pPB-TRE-DEST-rTA-HSV-neo (DOX>) (Villegas et al., 2019), respectively, and 1μg plasmid was reverse-transfected together with 1μg PBase and 3μL Lipofectamin 2000 in TbC1 cells. The next day fresh medium with the appropriate selection was added and the cells were analyzed after at least 1 week of selection.

### Immunofluorescence

Cells were plated on a laminin-coated 96-well glass plate (Greiner Bio-One), fixed with PBS-4% PFA for 20’, washed twice with PBS, permeabilized with PBS-0.1% Triton X-100 for 10’ at room temperature (RT), incubated in blocking solution (3% donkey serum (Sigma), 1% BSA in PBS-0.1% Tween-20 (PBST)) for 1h at RT, incubated with anti-Tfe3 antibody (Sigma, Cat# HPA023881, RRID:AB_1857931) 1/1000 in blocking solution overnight at 4°C, washed three times with PBST, stained for 2h at RT with secondary antibody Donkey Anti-Rabbit IgG – Alexa Fluor 555 (Life Technologies) 1/500 in PBST, counterstained with PBST-Hoechst33342 1/5000 (Life Technologies), washed twice with PBS and imaged at an LSM-710 scanning head confocal microscope (Zeiss). Images were exclusively cropped with no further manipulation.

### Western Blot

WT and Nmt1^-/-^ mESCs grown in 2iLIF were washed twice with PBS and incubated for 30’ or 6h in N2B27 + 12 ng/ml bFGF, put on ice, harvested, and lysed in RIPA buffer (50mM Tris pH7.5, 150mM NaCl, 1% IGEPAL, 0.5% Na Deoxycholate, 0.1% SDS, 2mM EDTA) with fresh Complete Protease Inhibitor Tablet (Roche) and Phosphatase Inhibitor Tablet (Roche). 10μg of cell lysate were resolved on a 10% SDS-PAGE and wet-blotted on nitrocellulose. Separate membranes were probed with Erk1/2 antibody (Cell Signaling Technology, #9102) or Phospho-Erk1/2 antibody (Cell Signaling Technology, #9101) 1/1000 in PBST-5% BSA.

### Affinity-purification mass-spectrometry

2*10^5^ WT or mutant naïve TNG-A cells were transfected in triplicate with 1μg pgk>BirA plasmid and 1μg DOX>prey plasmid according to Table S4 or no plasmid as negative control with 3μL Lipofectamin 2000. The next day fresh medium with hygromycin (selection for the pgk>BirA plasmid) was applied. Cells were grown in selective medium for 3-4 days and then the medium was changed to Serum-LIF + 1μg/mL Doxycycline. After 48 hours 10^7^ cells were harvested with trypsin, washed in PBS-0.1%BSA, washed in PBS, and nuclei were extracted in 10mM TrisHCl pH7.5, 10mM KCl, 1mM DTT, 0.5% IGEPAL, with Complete protease inhibitor (Roche) for 20’ minutes on ice. Nuclei were lysed by rotation for 1h at 4°C in 20mM TrisHCl 7.5, 100mMKCl, 1.5mM MgCl_2_, 1mM DTT, 10% glycerol, 0.5% TritonX-100, Complete protease inhibitor (Roche), Phosphatase inhibitor (Roche), and 250U/mL Benzonase (Merck). Lysates were clarified by centrifugation for 5’ at 12’000g at 4°C, 10μL of M280 Streptavidin-Dynabeads (ThermoFisher) were added and incubated at 4°C rotating for 4h. Beads were then washed three times with 20mM TrisHCl pH7.5, 150mM NaCl, with 0.5% IGEPAL and twice without IGEPAL. Beads were digested with Lys-C at RT for 4h and then with trypsin overnight at 37°C.

The generated peptides were acidified with TFA to a final concentration of 0.8% and analyzed by capillary liquid chromatography tandem mass spectrometry with an EASY-nLC 1000 using the two-column set-up (Thermo Scientific). The peptides were loaded with 0.1% formic acid, 2% acetonitrile in H_2_O onto a peptide trap (Acclaim PepMap 100, 75um x 2cm, C18, 3um, 100Å) at a constant pressure of 800 bar. Peptides were separated, at a flow rate of 150 nL/min with a linear gradient of 2–6% buffer B in buffer A in 3 minutes followed by an linear increase from 6 to 22% in 40 minutes, 22-28% in 9 min, 28-36% in 8min, 36-80% in 1 min and the column was finally washed for 14 min at 80% B (Buffer A: 0.1% formic acid, buffer B: 0.1% formic acid in acetonitrile) on a 50μm x 15cm ES801 C18, 2μm, 100Å column (Thermo Scientific) mounted on a DPV ion source (New Objective) connected to a Orbitrap Fusion (Thermo Scientific). The data were acquired using 120000 resolution for the peptide measurements in the Orbitrap and a top T (3s) method with HCD fragmentation for each precursor and fragment measurement in the ion trap according the recommendation of the manufacturer (Thermo Scientific). Protein identification and relative quantification of the proteins was done with MaxQuant version 1.5.3.8 using Andromeda as search engine (Cox et al., 2011) and label free quantification (LFQ, (Cox et al., 2014)) as described (Hubner et al., 2010). The mouse subset of the UniProt data base combined with the contaminant database from MaxQuant was searched and the protein and peptide FDR were set to 0.01.

The LFQ values were analyzed with Perseus v.1.6.2.2 as follows: entries identified only by site or reverse and potential contaminants were removed, values were Log2 transformed, entries identified in less than 2 replicates in any group were removed, and missing values were imputed based on the normal distribution of each replicate with a 0.25-fold width and a down-shift of 1.8-fold. Volcano plots are based on 2-sided t-test and threshold curves on an SO=0.1 and FDR=0.0054 for Zbtb2 baits and FDR=0.02 for Ubn1/2 baits.

For the heatmap representation in Figure 2B, missing Mascot values in experimental triplicates of Zbtb2 mutant AP-MS were imputed with a 1.8-fold downshift and a 0.25-fold distribution width of the actual distribution of detected proteins using the *fitdistr* function from the Cran package MASS (https://cran.r-project.org/web/packages/MASS/index.html), as described (Tyanova et al., 2016). To correct for varying Zbtb2 amounts in different APs, protein enrichments in each AP were normalized to the respective Zbtb2 bait. To compare interaction strengths, the enrichment of each interactor was normalized to its enrichment in wildtype Zbtb2 purifications.

### Yeast-2-hybrid

Yeast-2-hybrid assays were performed with the plasmids and strains from the Matchmaker Gold Yeast Two-Hybrid System (Takara Bio) according to manufacturer’s protocol, with the following plasmid modifications. For N-terminal AD-fusions, pGADT7 was digested with NdeI and BamHI and religated with the oligo TAGTGGTGGAACAAAAATGGGCCCGAATTCCCGGGATCGATTAACTGAGTAG, to create ApaI and ClaI sites for InFusion cloning (Takara Bio). For C-terminal -AD and -DBD fusions, TTTAAACTATTTGGGCCCATTTTTGTTCCACCACTATAAGCTTGGAGTTGATTGTATGCTTGG and either AAATGGGCCCAAATAGTTTAAACCGCGGTGGATCTGGTGGAATGGATAAAGCGGAATTAATTCCCG AG or AAATGGGCCCAAATAGTTTAAACCGCGGTGGATCTGGTGGAATGAAGCTACTGTCTTCTATCGAACA AGC were used for site-directed mutagenesis of pGADT7 and pGBKT7, respectively, to create a new MCS with ApaI and SacII sites for InFusion cloning (Takara Bio). pGBKT7-C and pGADT7-C were digested with NdeI and BamHI and religated with the oligo TATGCCAGCTGCTAAAAGAGTTAAATTGGATTAG to create a new c-Myc NLS. Briefly, pGBKT7 and pGADT7 plasmids were transformed into the yeast strains Y2HGold and Y187, respectively. Several colonies were picked and grown overnight in SD-Trp or SD-Leu, for pGBKT7 or pGADT7 plasmids respectively. When the cultures reached an OD600 ∼ 0.5, they were mated overnight in YPD medium and grown on an SC-Trp/-Leu plate for 2-3 days, and plate replicas were made on an SD-Ade/-His/-Trp/-Leu/+X-alpha-Gal/+Aureobasidin A plate. Pictures of the plates were taken after 2-4 days and images were exclusively cropped, with no further processing.

### Phylogeny

Btb domain sequences were retrieved with the ‘Architecture analysis’ tool of SMART (smart.embl.de; query: “BTB AND ZnF_C2H2” in “Mus musculus”) or individually with the ‘Sequence analysis’ tool. The phylogenetic tree was calculated, based on a multiple sequence alignment generated with T-Coffee (https://www.ebi.ac.uk/Tools/msa/tcoffee/), using the neighbour-joining clustering method provided by the ClustalW2 package (http://www.clustal.org/clustal2/) running 1000 iterations. The tree was visualized with iTOL (https://itol.embl.de/).

### Gene expression analysis

RNAseq reads and published data (see table below) were aligned to the mouse GRCm38/mm10 genome using *qAlign* from the Bioconductor package QuasR (Gaidatzis et al., 2015) with default parameters except for *aligner=“Rhisat2”* and *splicedAlignment=TRUE*. For aligning RNAseq reads for Dux overexpression (Hendrickson et al., 2017), *paired=“fr”* was additionally used. Alignments were quantified for known UCSC genes obtained from the TxDb.Mmusculus.UCSC.mm10.knownGene package using *qCount*. Microarray data (Fazzio et al., 2008) was analyzed and normalized using the Bioconductor package limma (Ritchie et al., 2015). In this dataset, Tip60 knockdown replicate 3 is an outlier, and Ep400 knockdown replicates 1 and 3 do not show Ep400 transcript reduction, and were therefore excluded from the analysis.

Differential gene expression (Table S3) was determined using edgeR (Robinson and Oshlack, 2010). In Fig. 1G, Fig. S1G, and Fig. S5A preEPI is the combination of the EPI and ICM+EPI markers, and postEPI the PE geneset (Boroviak et al., 2015). Pearson correlation coefficients (Fig. 5C) were calculated using R’s *cor* function. For heatmap visualization (Fig. 5D), significantly deregulated genes in Zbtb2 mutants were considered (Table S3), which are the genes that showed an absolute log2 expression FC of greater than 1 with a false discovery rate of less than 0.005 between wildtype and Zbtb2 KO cells in at least one of the conditions: 2iLIF, 24h EpiLC, 48h EpiLC, 24h Serum-LIF, 48h Serum-LIF. All contrast shown in Fig. 5D were used for clustering.

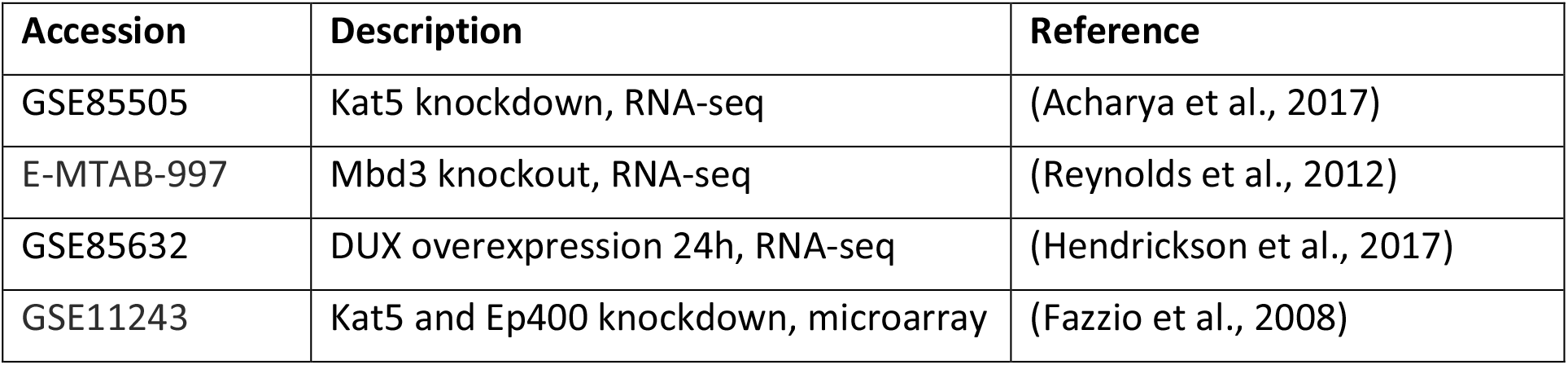

### ChIPseq analysis

Published datasets (see table below) were aligned to the mouse GRCm38/mm10 genome using *qAlign* and profiled using *qProfile* from the Bioconductor package QuasR. 137435 ATAC peaks (Olivieri et al., 2020) were called using Macs2 (Zhang et al., 2008), of which 22826 were in promoters (+/- 1kb of annotated transcriptional start sites). For heatmap visualization (Fig. S5D), ChIPseq signals were profiled in these promoters and ChIP enrichment calculated over respective inputs (Zbtb2, H3K4me3, Chd4, Mbd3, Dux) or controls (Kat5, Ep400). For metaplots (Fig. 5E, F, Fig. S5B, C), ChIPseq signals were profiled in promoter regions of cluster genes that were extracted from the TxDb.Mmusculus.UCSC.mm10.knownGene Bioconductor package (https://bioconductor.org/packages/release/data/annotation/html/TxDb.Mmusculus.UCSC.mm10.knownGene.html).

| Accession | Description | Reference |
| --- | --- | --- |
| GSE85632 | DUX 18h, ChIP-seq | (Hendrickson et al., 2017) |
| GSE101802 | Zbtb2 2i, ChIP-seq | (Karemaker and Vermeulen, 2018) |
| E-MTAB-6804 | Mbd3 and Chd4, ChIP-seq | (Bornelöv et al., 2018) |
| GSE67584 | Kat5 and Ep400, ChIP-seq | (Chen et al., 2015) |
| GSE74112 | K3K4me3, ChIP-seq | (Liu and Kraus, 2017) |
| E-MTAB-9453 | ATAC-seq | (Olivieri et al., 2020) |

### RT-qPCR

RNA was extracted from naïve mESCs with the RNeasy Mini Kit (Qiagen) and 1μg of it was reverse-transcribed with SuperScript III Reverse Transcriptase (LifeTechnologies). qPCR was performed with the TaqMan Fast Universal PCR Master Mix (ThermoFisher) and the TaqMan probes GAPDH (4352339E) and Zbtb2 (Mm01605943_g1). The DOX>Zbtb2 plasmid that cannot be detected by Mm01605943_g1 was produced by fusion PCR of the endogenous cDNA and the following gBlock (Integrated DNA Technologies): ATGGATTTGACCAACCATGGACTTATTCTACTGCAGCAGTTAAACGCTCAGCGAGAGTTTGGTTTCCTGTGTGACTGCACGGTTGCAATCGGCGATGTGTATTTTAAAGCCCATAAGAGTGTGTTGGCAAGTTTTAGTAACTATTTCAAAATGCTTTTCGTGCACCAAACATCAGAGTGTGTGAGATTAAAACCAACAGATATCCAACCAGATATCTTTTCATACTTATTGCATTTAATGTATACCGGGAAGATGGCCCCACAGCTCATCGACCCTGTGAGGCTAGAGCAAGGGATCAAATTCCTGCACGCATACCCCCTCATCCAGGAAGCCAGCCTTGCCAGCCAAGGCAGCTTTTCCCATCCCGAGCAAGTCTTCCCTCTGGCCTCATCCTTGTACGGCATTCAGATTGCAGACCATCAGCTGAGACAAGCCACCAAGATGAATTTAGGGCCTGAGAAACTTGGACGGGAGCCTAGGCCACAGGCATCCAGGATGA. The construct was validated by plasmid qPCR with Mm01605943_g1.

### Data availability

The genomic data generated for this study have been deposited at ArrayExpress (E-MTAB-9796, E-MTAB-9797, E-MTAB-9798). The mass spectrometry proteomics data have been deposited at the ProteomeXchange Consortium via the PRIDE partner repository with the dataset identifiers PXD022451 and PXD022446.

## FIGURES

**Supplementary Figure S1:**
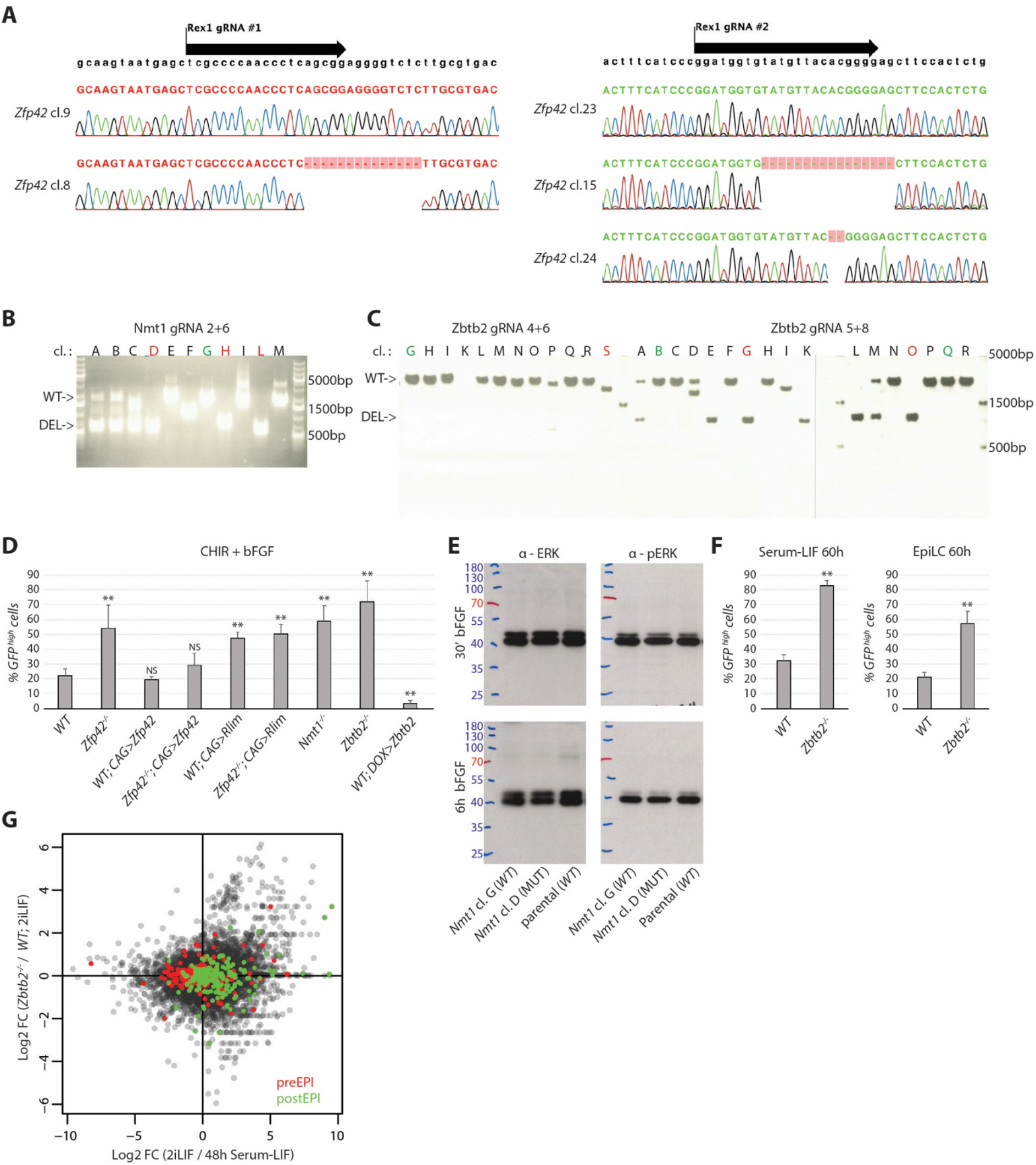
Related to Figure 1. **(A**) Chromatograms of *Zfp42*^-/-^ and WT sibling clones. (**B**,**C**) Genotyping PCR of *Nmt1* and *Zbtb2* knockout and sibling control clones used in this study. **(D**,**F)** Average and standard deviation (SD) of *Nanog>GFP*-high cells of biological triplicates quantified as in Fig.1B, C, D (**D**). and Fig.1F (**F**). ** indicates p-values<0.001 and NS p-values>0.1 compared to corresponding *WT* controls. **(E)** Anti-ERK and anti-phospho-ERK western-blot of lysates from *Nmt1*^-/-^ and *WT* clones. **(G)** Scatterplot of log2FCs in gene expression of indicated contrasts. Green labels post-implantation and red pre-implantation epiblast specific genes (Boroviak et al., 2015).

**Supplementary Figure S2:**
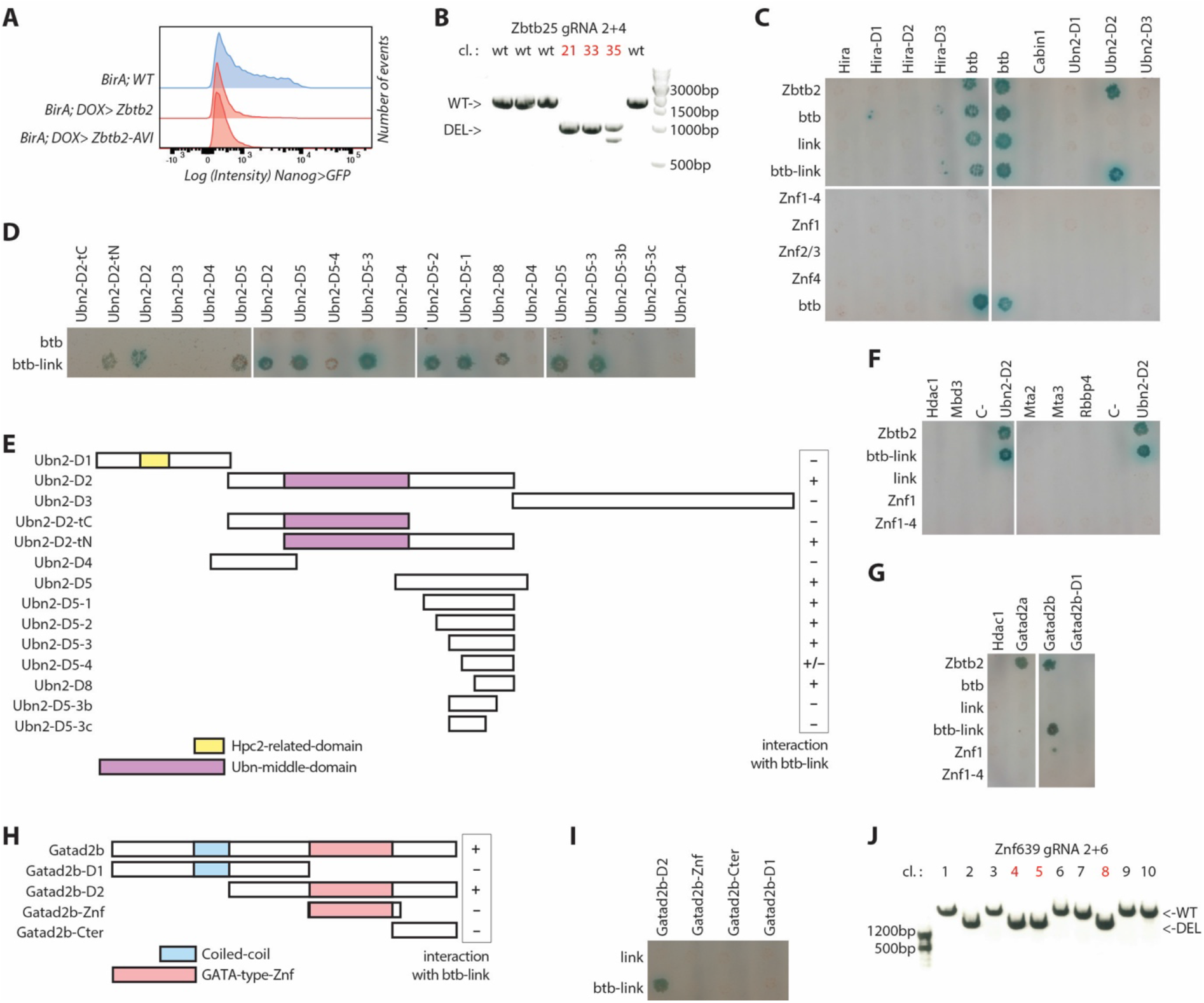
Identification of the ZBTB2-interacting subunits and domains. Related to Figure 2. **(A)** *Nanog>GFP* intensities upon ZBTB2 or ZBTB2-AVI induction for 3 days in N2B27 +CHIR +bFGF. **(B)** *Zbtb25* genotyping PCR of *Zbtb2*^-/-^; *Zbtb25*^-/-^ clones; red labels clones used in this study. **(C**,**D**,**F**,**G**,**I)** As in Fig. 2D (**C**,**F**,**G**) and Fig. 2E (**D**,**I**). **(E**,**H)** Diagram of *Ubn2* (**E**) and *Gatad2b* (**H**) constructs used for Y2H analysis. **(J)** Genotyping PCR of *Znf639* mutant clones; red labels clones used in this study.

**Supplementary Figure S3:**
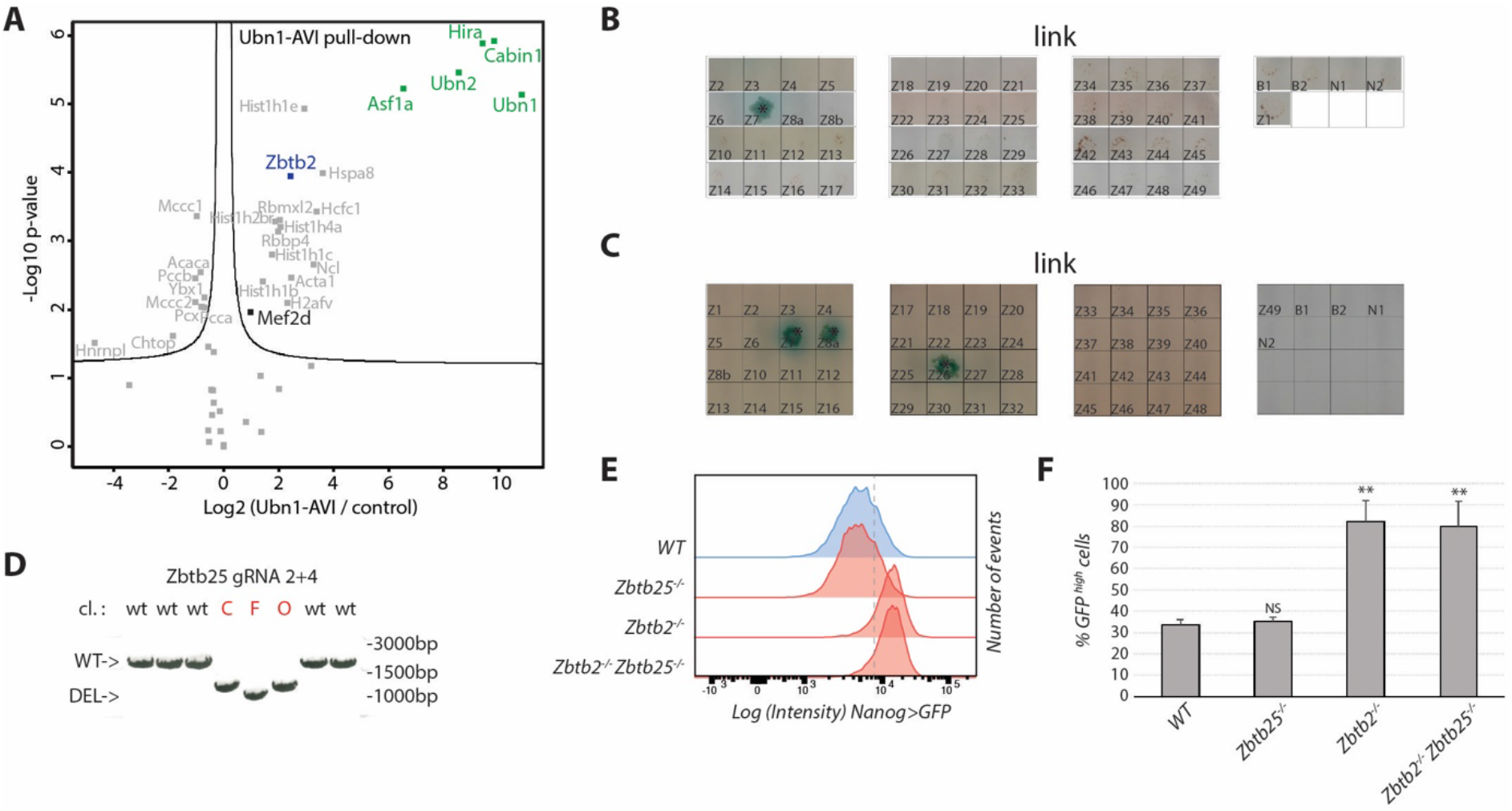
Lack of genetic interaction between ZBTB2 and ZBTB25. Related to Figure 3. **(A)** Volcano plot of protein enrichments in AP-MS of UBN1-AVI compared to control BirA-expressing cells in 2iLIF. ZBTB2 is indicated in blue and HiRA subunits in green. (**B**,**C**) Colony growth of control matings for experiments presented in Fig.3B,C,F **(B)** and in Fig.3D **(C)** using ZBTB2’s link region as control bait construct. **(D)** *Zbtb25* genotyping PCR of *Zbtb25*^-/-^ clones; red labels clones used in this study. **(E**,**F)** *Nanog>GFP* intensities of indicated genotypes after 60h in Serum-LIF (**E**). Dashed line indicates the threshold for quantification of GFP-high cells as average and SD of biological triplicates **(F)** ** indicates p-values<0.001 and NS p-values>0.1 compared to the *WT* control.

**Supplementary Figure S4:**
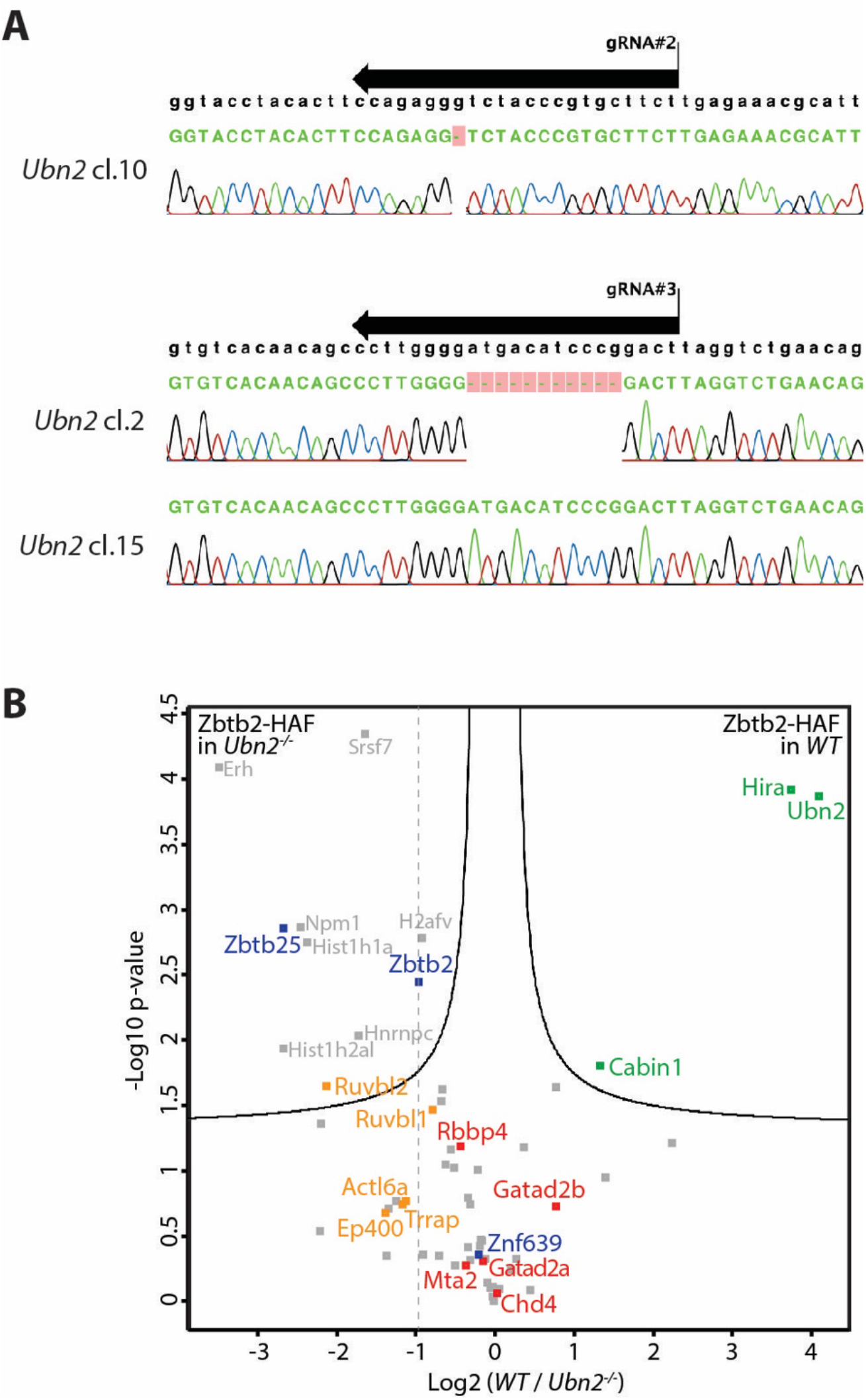
The Ep400 interaction is not mediated by the HiRA complex. Related to Figure 4. **(A)** Chromatograms of *Ubn2*^-/-^ and sibling *WT* clones. **(B)** Volcano plot of protein enrichments in AP-MS of ZBTB2-HAF in *WT* compared to *Ubn2*^-/-^cells. ZBTB2 and partner TFs are indicated in blue. The dashed line marks enrichment of the ZBTB2-HAF bait protein which is similar to that of NuRD subunits in red and Ep400 subunits in orange, while interaction with HiRA subunits in green is comparatively reduced in *Ubn2* mutant cells.

**Supplementary Figure S5:**
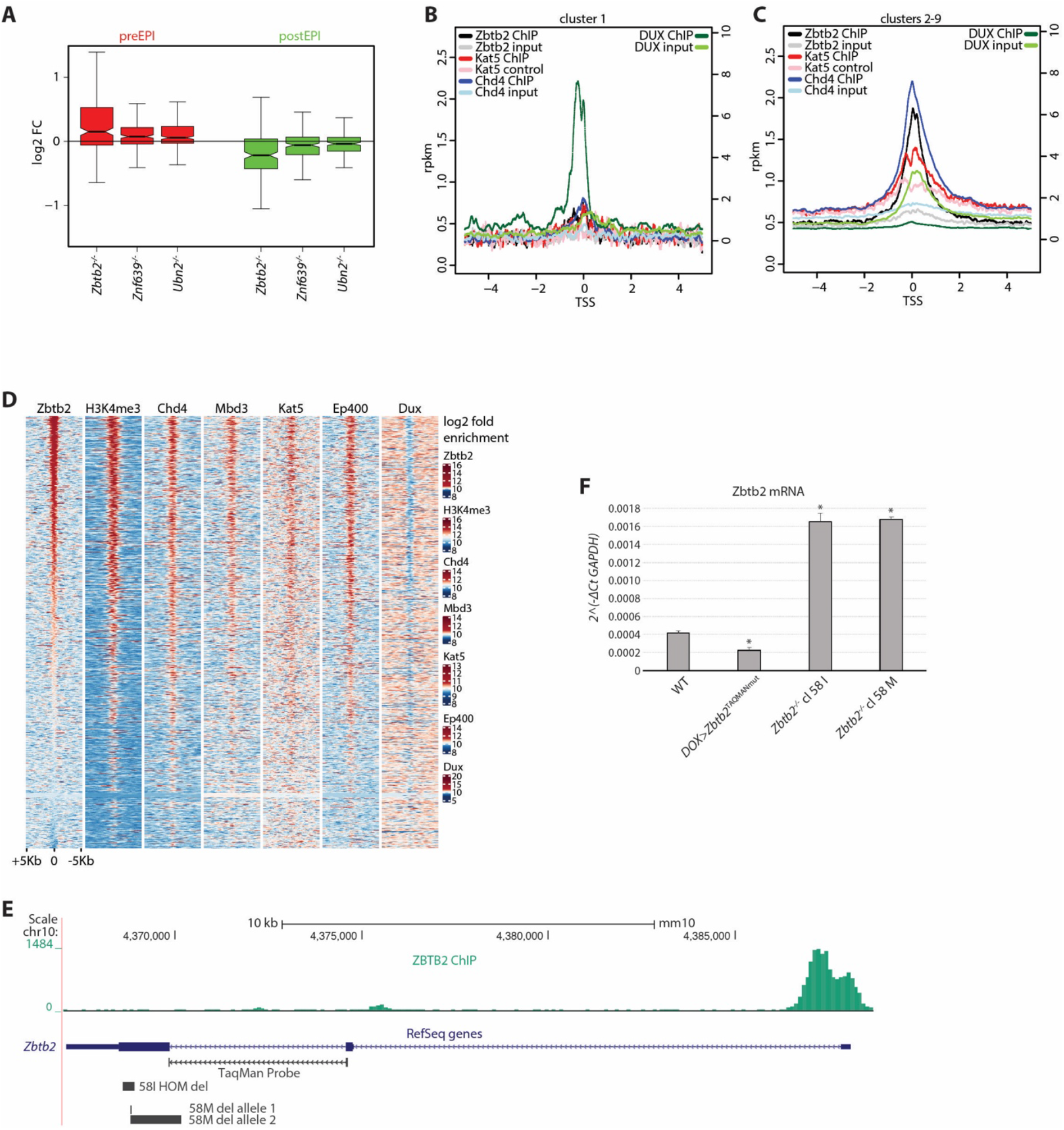
ZBTB2 binds and represses its own promoter. Related to Figure 5. **(A)** Boxplot of log2 fold differential expression after 48h in Serum-LIF of post-implantation and pre-implantation epiblast specific genes (Boroviak et al., 2015) in indicated genotypes compared to *WT* cells. **(B**,**C)** ZBTB2 (Karemaker and Vermeulen, 2018), KAT5 (Chen et al., 2015) and CHD4 (Bornelöv et al., 2018) (left scale), and DUX (Hendrickson et al., 2017) (right scale) ChIP-seq rpkm centered on TSSs of cluster 1 (**B**) and cluster 2-9 genes (**C**). **(D)** Heatmap of ZBTB2 (Karemaker and Vermeulen, 2018), H3K4me3 (Liu and Kraus, 2017), CHD4, MBD3 (Bornelöv et al., 2018), KAT5, EP400 (Chen et al., 2015) and DUX (Hendrickson et al., 2017) log2 fold ChIP-seq enrichment over respective controls at accessible (ATACseq, not shown) TSSs. **(E)** Diagram of the *Zbtb2* locus, showing the ZBTB2 peak at the TSS, and the deletions in the *Zbtb2*^-/-^ clones 58I and 58M that give rise to transcripts that are detectable by qPCR probes against endogenous *Zbtb2*. **(F)** Average and SD of technical triplicates of endogenous *Zbtb2* relative to *GAPDH* transcription in *WT* cells, *WT* cells over-expressing a *Zbtb2* construct that cannot be detected by the qPCR probe (TAQMANmut) and the *Zbtb2*^-/-^ clones depicted in Fig. S2E. * indicates p-values<0.01 relative compared to *WT*.

## SUPPPLEMENTARY TABLES

**Supplementary Table S1: Screen results**

**Supplementary Table S2: CRISPR KO and genotyping strategies; cell lines**

**Supplementary Table S3: Gene expression tables with clusters and post- and pre-implantation genes**

**Supplementary Table S4: AP-MS bait sequences and Y2H constructs**

**Supplementary Table S5: AP-MS results tables**

